# Counterfactual Diffusion Models for Interpretable Explanations of Artificial Intelligence Models in Pathology

**DOI:** 10.1101/2024.10.29.620913

**Authors:** Laura Žigutytė, Tim Lenz, Tianyu Han, Katherine Jane Hewitt, Nic Gabriel Reitsam, Sebastian Foersch, Zunamys Itzell Carrero, Michaela Unger, Asier Rabasco Meneghetti, Alexander T. Pearson, Daniel Truhn, Jakob Nikolas Kather

## Abstract

Deep learning can extract predictive and prognostic biomarkers from histopathology whole slide images. However, explainable artificial intelligence approaches widely used in digital pathology, such as attention heatmaps and class activation mapping, offer only limited interpretability regarding the features captured by classifiers. Here, we present MoPaDi (Morphing histoPathology Diffusion), a framework for generating counterfactual explanations for histopathology images that reveal which morphological or style features drive classifier predictions. MoPaDi combines diffusion autoencoders with task-specific multiple instance learning classifiers to manipulate images and flip predictions by modifying relevant features. We evaluated the framework on multiple datasets spanning colorectal, breast, liver, and lung cancers, including tissue type, cancer subtype, and biomarker (microsatellite instability) classification tasks. We assessed counterfactual explanations through quantitative analyses, pathologists’ evaluations, and independent foundation model-based classifiers. We found that MoPaDi was able to generate realistic counterfactual histopathology images, enabling pathologists to identify morphological features associated with the change in model predictions. Unlike conventional reviews of highly attended regions typical in digital pathology, MoPaDi explanations enabled pathologists to directly identify morphological features driving the classifier’s predictions from a limited number of top-contributing tiles. Consistent with the literature, our biomarker classifier associated high microsatellite instability with mucinous differentiation, glandular patterns, and lymphocytic infiltration. Furthermore, MoPaDi revealed that changes in classifier predictions were mainly driven by morphological alterations rather than staining differences. Overall, MoPaDi is a practical framework for counterfactual explanations in computational pathology that reveals model-specific drivers of classification and increases trust in deep learning models.

## Introduction

Deep learning (DL) can predict diagnostic and prognostic biomarkers from hematoxylin and eosin (H&E) stained whole slide images (WSIs) derived from pathology slides of cancer (1–4). DL performs well for many clinically relevant tasks, such as automating pathology workflows to make diagnoses or extract subtle visual information related to biomarkers (5). However, the so-called ‘black box’ nature of DL can be problematic because it is often unclear which visual features the model bases its predictions on, limiting interpretability and trust in the classifier (6). Established explainability methods, such as feature visualization (7) and pixel attribution (e.g., saliency maps, including attention heatmaps) (8), provide limited insight and cannot ensure that model decisions arise from meaningful or generalizable morphological features rather than dataset-specific artifacts (6). Although widely used in biomedical research, interpreting these methods is challenging and can result in overestimating the model’s performance due to confirmation bias (6,9,10).

An alternative approach, counterfactual explanations (11,12), has been explored in other domains, such as natural images, radiology, and dermatology (13–15), but is underexplored in computational pathology. Counterfactual explanations ask ‘What if?’ questions, such as ‘How would an image need to change to be classified as class X?’ (**Fig. 1A**). Here, we developed and validated MoPaDi (Morphing histoPathology Diffusion), a framework that generates classifier-guided counterfactual edits for histopathology images. Images generated in this way show features (morphological or style-related, e.g., staining) that the DL classifier is sensitive to when making predictions. The objective is to make the decision-making process of ‘black-box’ artificial intelligence (AI) models more understandable and trustworthy.

**Figure 1.**
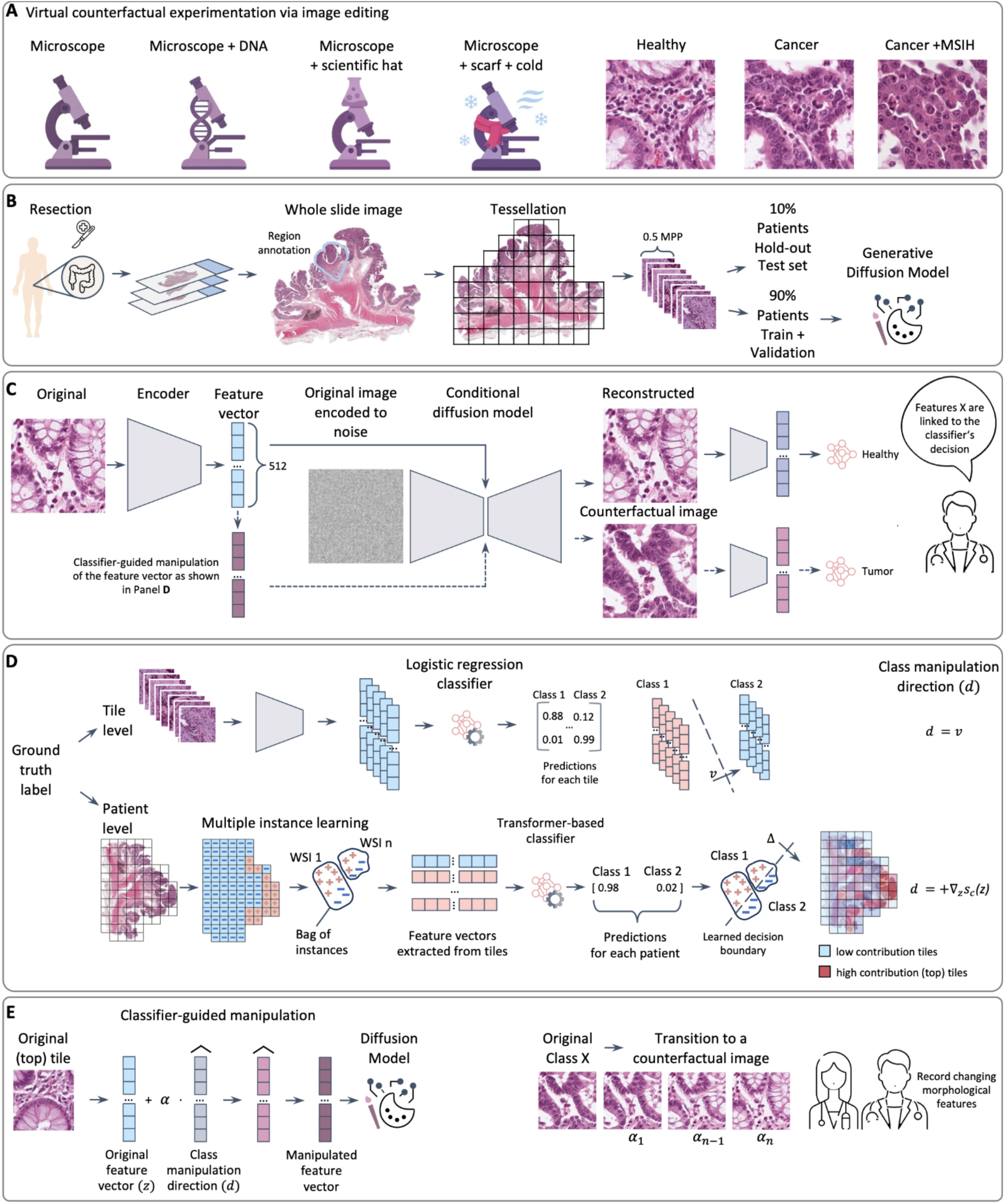
Overview of the experimental setup. (**A**) Conceptual illustration showing how semantic modifications to an input can generate counterfactual variants. Analogous to adding attributes to a microscope (**left**), our method alters key biological attributes in tissue images (**right**), demonstrating transitions from healthy morphology to cancer and then cancer with microsatellite instability-high features. ‘What-if’ (counterfactual) edit denotes a model-internal, decision-linked transformation, while not implying biological causality. (**B**) Preprocessing steps. Digitized whole slide images (WSIs) were resized to 0.5 micrometers per pixel (MPP), tessellated into 512×512 px tiles, and filtered to include only pathologist-annotated tumor regions. (**C**) MoPaDi is trained simultaneously as a feature extractor and a denoising diffusion implicit model (DDIM), which is conditioned on feature vectors and the original image encoded into noise, enabling the reconstruction of original images. (**D**) When a ground truth label is available on a tile level (NCT-CRC-HE-100K dataset), a logistic regression classifier is trained on tile-level feature vectors extracted with MoPaDi encoder to define the decision boundary used for classifier-guided counterfactual generation. Binary classification is shown here for illustrative purposes; the method generalizes to multiclass settings using a one-vs-rest approach. Meanwhile, when a ground truth label is available on a patient level, a multiple instance learning (MIL) approach is used. MoPaDi learns patient-level predictions from bags of tile features using a transformer-based classifier with attention-based pooling (refer to **Fig. S1** for more extensive transformer architecture details). (**E**) For both classifier types, counterfactual images are generated by manipulating the feature vector of a selected tile with a manipulation amplitude *α*. In tile-level classification, *d* follows the classifier weight vector 𝑣; in the MIL case, *d* is computed as the gradient of the classifier’s output *s_c_(z)* for a chosen counterfactual class *c* with respect to the tile’s feature vector *z*. The denormalized manipulated feature vector is then used to condition DDIM, yielding a counterfactual image. Pathologists were shown transitions from original to counterfactual images and asked to identify key morphological changes.

Previous studies in other image analysis domains, such as natural images or radiology, have shown that counterfactual explanations can help uncover biases and confounding variables (14,16), as well as provide model prediction explanations. However, these efforts have largely been limited to radiology data or outdated generative AI models such as generative adversarial networks (GANs) (14,17,18). Although GAN-based methods have been explored previously (15,19–24), they are less stable during training and produce lower-quality synthetic samples. Therefore, we propose utilizing state-of-the-art diffusion models to generate counterfactual explanations for digital pathology. Diffusion models have surpassed GANs in both synthetic image quality and training stability (25). They work by gradually transforming images into noise and learning the reverse process, thereby enabling the generation of realistic images from the learned data distribution (26). A key strength of diffusion models over GAN-based methods is their ability to modify real images while preserving details unrelated to the specific manipulation (27), due to the iterative nature of their image transformation process, which allows fine-grained control over which features are transformed within an image (28). Furthermore, most computational pathology problems require a weakly supervised approach since labels are typically available only for WSIs consisting of many tiles (**Fig. 1B**), but not for every tile individually (29). These tasks are usually addressed by multiple instance learning (MIL) (30). Therefore, we extend the concept of counterfactual image generation to the weakly supervised scenario within the present study, in which we train an attention-based classifier on bags of tiles with a patient-level label, and then generate counterfactual explanations for the most predictive tiles. MoPaDi combines diffusion autoencoders (27) with MIL to generate high-quality counterfactual images of histopathology tiles, preserving original image details and enabling pathologists to comprehensively evaluate morphologies captured by the classifier (**Fig. 1C–E**). To the best of our knowledge, diffusion-based explainability methods have not been applied to histopathology data for counterfactual image generation combined with MIL.

The aim of MoPaDi is to reveal which morphological features in histopathology images drive the predictions of DL classifiers by creating counterfactual explanations. To validate our framework, we first show that the model learns histology-relevant morphological features by generating tissue-type targeted edits. We then extensively evaluate MoPaDi for exposing discriminative features of a microsatellite instability (MSI) status classifier, a colorectal cancer (CRC) biomarker whose predictability from H&E-stained WSIs has been widely demonstrated (3,31). We also show how counterfactual explanations can be used to disentangle model predictions into stain-style and morphology-related components, and quantitatively demonstrate that class shifts for MSI predictions are predominantly driven by morphology. Furthermore, we validate MoPaDi on pan-cancer, clinically relevant tasks, including the classification of hepatobiliary, lung, and breast cancer types. Our framework, validated across diverse datasets, reveals key morphological features driving the classifier’s predictions through automated, targeted manipulation of histopathological images toward counterfactual images, enabling understanding of morphological patterns that the DL model is sensitive to.

## Materials and Methods

### Experimental Design

We designed a multi-stage evaluation to (i) learn a faithful, decodable latent representation of histopathology images, (ii) train task-specific classifiers at the tile and patient level, and (iii) generate and validate classifier-guided counterfactual images that reveal decision-relevant morphology (**Fig. 1**).

First, MoPaDi diffusion autoencoders were trained on multiple datasets to enable high-fidelity reconstructions and controllable editing: NCT-CRC-HE-100K, The Cancer Genome Atlas Research Network (TCGA) colorectal cancer cohort (TCGA-CRC), breast cancer cohort (TCGA-BRCA), and a TCGA pan-cancer collection of histopathology tiles. Reconstruction fidelity and generative realism were quantified on test sets using standard quantitative metrics. We then established two classification approaches from MoPaDi’s encoder’s features (**Fig. 1D**): (a) a tile-level logistic regression (one-vs-rest) for multiclass tissue typing in colorectal cancer, and a weakly supervised, transformer-based MIL model for slide/patient-level tasks.

We then generated counterfactual images to expose what morphological changes the classifier is sensitive to by producing visual, model-consistent feature shifts sufficient to alter predictions. We validated effectiveness by summarizing the fraction of tiles whose predicted class flipped to the target class, confirming trends with independent classifiers, disentangling staining (style) from structure (morphology), and quantifying cellular and structural changes along counterfactual transitions. To assess biological plausibility and perceptual realism, two expert studies were conducted: (a) pathologists annotated original to counterfactual transitions for emerging/disappearing histologic features, and (b) a blinded real-vs-synthetic study measured perceptual realism.

Together, this design assesses (1) diffusion autoencoding fidelity for histopathology, (2) classifier performance, (3) counterfactual effectiveness, (4) expert interpretability and perceptual realism, (5) independence from stain/style, and (6) quantitative cellular correlates – across multiple cancers and tasks.

### Datasets and Preprocessing

This study explored two colorectal cancer datasets, starting with the publicly accessible NCT-CRC-HE-100K dataset (32). This dataset comprises 100,000 H&E-stained histopathological images obtained from 86 formalin-fixed paraffin-embedded samples representing both human colorectal cancer and normal tissues. The dataset encompasses nine distinct tissue classes: adipose, background, debris, lymphocytes, mucus, smooth muscle, normal colon mucosa, cancer-associated stroma, and colorectal adenocarcinoma epithelium (**Fig. S5A**). All classes were used to train the diffusion auto-encoder. Each image had a resolution of 224 x 224 pixels at 0.5 micrometers per pixel (MPP), and was color-normalized using Macenko’s method (33). The trained model was evaluated on 7,180 tiles from the CRC-VAL-HE-7K dataset, comprising 50 WSIs, which was published alongside NCT-CRC-HE-100K, and serves as a non-overlapping dataset for testing purposes (32). Further analysis was performed on a second colorectal cancer dataset from TCGA, consisting of 625 WSIs obtained from formalin-fixed paraffin-embedded samples. This cohort was randomly divided at the patient level, allocating 90% for training and 10% for independent testing before any preprocessing. The held-out test set was used not only to evaluate classifier performance but also to assess the quality of image reconstruction, generation, and manipulation. For the TCGA-CRC dataset, in which raw whole-slide images were processed directly, the slides were tessellated into tiles of 512 x 512 pixels at a spatial resolution of 0.5 MPP. WSIs lacking original MPP metadata (*N* = 26) or tumor annotations (*N* = 8) were excluded, resulting in a set of 588 WSIs from 582 colorectal cancer patients. Patient characteristics and counts per split are provided in **Table S1**. Tiles were derived from tissue regions determined by Canny edge detection. Tiles derived from non-tumorous regions were filtered out, retaining only those originating from tumor regions. This selection process was guided by coarse tumor area annotations prepared by pathology trainees and verified by an expert pathologist. Tiles overlapping by at least 60% with annotated tumor regions were included. No color normalization was performed (except for the NCT-CRC-HE-100K dataset, which was pre-normalized) to facilitate the investigation of staining influence on classifiers. To determine the threshold between MSI (MSIL) and MSI-high (MSIH), we fitted a two-component Gaussian mixture model to the log-transformed distribution of total MSI events (obtained from Cortes-Ciriano et al. (34)) across TCGA-CRC samples and defined the cutoff (*τ*) at the intersection of the two components, with microsatellite stable (MSS) = 0, MSIL < *τ*, and MSIH ≥ *τ*.

Further investigation was conducted on the TCGA-BRCA cohort (1,115 WSIs from 1,046 patients) to investigate counterfactual explanations for breast cancer. This dataset was preprocessed using the same pipeline applied to TCGA-CRC. Finally, we used another publicly available dataset, further referred to as the TCGA Pan-cancer dataset, where varying resolution tiles (256 x 256 pixels at MPP ranging from 0.5 to 1) were extracted from tumor regions from WSIs of 32 cancer types (35). Patient counts in test and train splits are provided in **Table S2**.

### Diffusion Autoencoders

MoPaDi’s architecture builds on the official diffusion autoencoders implementation by Preechakul et al. (27). It consists of a denoising diffusion probabilistic model (DDPM; *D_θ_*), conditioned on a feature vector obtained with a learnable convolutional neural network encoder (*E_ψ_*). We refer to this encoder as the feature extractor, which learns to extract meaningful high-level information (feature vectors *z*) from input images (**Fig. 1C**). The encoder and decoder are trained jointly by optimizing a simplified noise prediction loss:

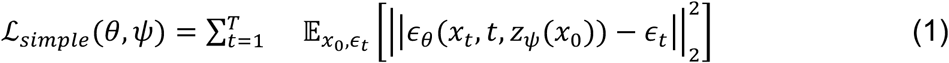

where ɛ*_θ_* is the predicted noise at diffusion timestep *t*, *x_t_* is the noisy image at that step, *z* is the encoded feature vector, and ɛ*_t_ ∼*𝒩(0,I) is the sampled Gaussian noise. Both *θ* and *ψ* are optimized jointly, encouraging the encoder to learn task-agnostic features while enabling high-fidelity decoding.

For training the autoencoding MoPaDi model on the NCT-CRC-HE-100K dataset, we used 8 Nvidia A100 GPUs with 40 GB of memory each. For the autoencoding model trained on TCGA-CRC, which was tessellated into larger tiles (512 x 512 px), we used 4 NVIDIA A100 GPUs with 80 GB of memory each. The models were trained using the DDPM objective, with denoising diffusion implicit model (DDIM) sampling applied at inference for efficient generation. Models were trained until an approximate Fréchet Inception Distance (FID) score below 20 was achieved (fast evaluation on 5,000 images). A global batch size of 24 for 512 x 512 px models and 64 for smaller models, divided evenly among the GPUs, was used. Models were optimized with Adam (learning rate 10^-4^ and no weight decay) with gradient clipping and exponential moving average of weights (decay 0.9999); no learning-rate scheduler was used. To evaluate the quality of the latent space learned by the autoencoding model and generate novel synthetic images, we additionally train a latent DDPM on the distribution of feature vectors obtained from the diffusion-based encoder.

### Classifiers and Counterfactual Image Generation

To enable classifier-guided counterfactual image generation, we first establish classification models for tile-level and patient-level ground truth labels. For tile-level tasks, such as tissue type classification, we train a multiclass logistic regression classifier on 512-dimensional features extracted with a pretrained MoPaDi feature extractor (**Fig. 1D**). Each tile is annotated with a one-hot target vector indicating its true class. Features are normalized to zero mean and unit variance before classification. Although the task is multiclass, we treat it as a one-vs-rest problem by applying binary cross-entropy loss independently to each class. The training process involved 300,000 tile exposures.

For patient-level ground truth labels, such as biomarker status, e.g., MSI, we adopt a MIL approach using transformer-based pooling. A transformer-based classifier processes bags of tile-level features, applying layer normalization, multi-head cross-attention, and a self-attention block, followed by a classifier head with two fully connected layers (with dropout and SiLU activation (36)), and a final linear output layer (**Fig. S1**). The classifiers were trained using the Adam optimizer, a learning rate of 0.0001, early stopping with a patience of 20 epochs, and cross-entropy loss with inverse class proportion weighting to address class imbalance. Model variability was assessed via 5-fold cross-validation. For counterfactual image generation, a final classifier is trained on the full training set, using 50–200 epochs based on the cross-validation results.

Both the linear and MIL classifiers guide counterfactual image generation by defining a class-specific direction of manipulation in feature space. In the linear case, the learned weight vector 𝑣 of the logistic regression model defines the direction *f* in feature space that increases the logit for the target class. This direction is used to shift the tile’s feature vector across the decision boundary. In the MIL setting, where the relationship between individual tiles and the label is non-linear, we compute the gradient of the classifier’s output for a chosen “counterfactual” class *c* with respect to the selected tile’s normalized feature vector *z* and use this gradient as the direction of manipulation, where *s_c_*(*z*) denotes the raw logit that the classifier assigns to a class 𝑐 given a feature vector *z*:

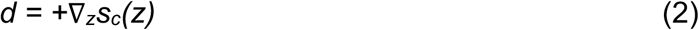

After modifying the original feature vector 𝑧 by adding *d* (scaled by the manipulation amplitude α and adjusted for feature dimensionality, i.e., 512 for MoPaDi encoders *E*) (**Eq. 3**), the updated feature vector *z* is de-normalized and passed to the conditional diffusion decoder (*D*), along with the original image encoded to noise (*x_T_*), to generate the counterfactual image *x^* (**Fig. 1E**, **Eq. 4)**.

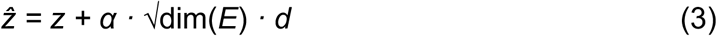

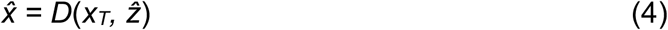

We apply counterfactual manipulation by progressively increasing α along the target-class gradient direction in latent space. For each dataset, we use a fixed set of manipulation amplitudes (e.g., α ∈ {0.2, 0.4, 0.6, 0.8, 1.0} for the linear approach and α ∈ {0.02, 0.04, 0.06, 0.08} for the MIL approach). Even if some manipulated tiles do not cross the classifier’s decision boundary at the largest *α*, we apply the same α values uniformly across the cohort to ensure comparability. We increase α until the majority of re-encoded counterfactual images are classified as the target class. The predictions are based on what the classifier would predict if it only had seen the particular tile. For the MIL approach, we generate counterfactual images for the top 5 most contributing tiles per WSI, with each tile being manipulated individually.

### Evaluation of Classifiers and Reconstructions

We assessed the autoencoder’s reconstruction quality on a randomly selected set of 1,000 images per dataset using the mean square error (MSE) between the original and reconstructed pixel intensities and multi-scale structural similarity index measure (MS-SSIM) (37). In addition, the structural similarity index (38) was used to visualize regions of similarity and dissimilarity between original and reconstructed images, following conversion to grayscale to facilitate interpretation. The latent space’s ability to retain semantically meaningful information was further evaluated by training an additional DDIM on feature vectors extracted by the MoPaDi encoder and sampling 10,000 synthetic images. To compare the distribution of real and generated images, we computed the FID score using the clean-fid library (39), with lower scores indicating greater similarity between real and synthetic image distributions. For comparing reconstructions with a GAN-based approach, we applied HistoXGAN (24) by resizing each tile (224 × 224), normalizing, extracting CTransPath features, and feeding them into the pre-trained StyleGAN3 generator to synthesize the reconstructed image.

To evaluate classifier performance, we computed the area under the receiver operating characteristic (ROC) curve on the held-out test set. Bootstrapping was employed as a resampling technique to estimate robustness and confidence intervals (CI). Resampling was done with replacement 1,000 times. We constructed ROC curves for each resampled set and interpolated between them to obtain a set of smoothed curves. 95% CI were extracted from smoothed curves. The latent space and the direction of manipulation were visualized using t-SNE, enabling a qualitative representation of extracted 512-dimensional feature vectors in a reduced two-dimensional format.

### Style-Morphology Decomposition of Prediction Changes

To quantify the relative influence of stain style versus tissue morphology on model predictions, we performed a Shapley-based style-morphology decomposition using pairs of original and counterfactual images. For each tile *x* (original) and its counterfactual *x_cf_* generated by the MoPaDi diffusion model, we defined the total prediction change for the target class *c* as:

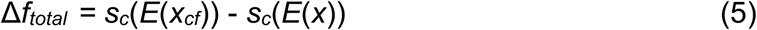

where *s_c_*(*·*) denotes the raw classifier logit.

To separate this change into style- and morphology-related components, we created two hybrid images:

● *x_style_* – preserving the morphology of the original tile while adopting the stain style of the counterfactual;
● *x_morph_* – preserving the stain style of the original while adopting the morphology of the counterfactual.

Stain style transfer was performed using the Vahadane method (40). Predictions on these hybrids were used to compute first-order effects of style and morphology. Following a Shapley-style additive decomposition, the total change was partitioned into:

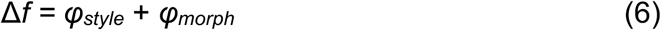

where *φ_style_* and *φ_morph_* represent the respective contributions of stain style and tissue morphology to the overall logit change, including half of their shared interaction term so that the sum equals the total change.

Predictions were obtained from the same MIL classifier used in the main experiments. The decomposition was applied per tile, and per-patient statistics were computed as the median of tile-level values. This approach quantifies how much of the model’s decision change arises from staining differences versus morphological differences.

### User Studies

To evaluate whether MoPaDi-based classifiers capture meaningful morphology, we conducted three complementary expert studies across tasks. First, to assess whether the classifiers capture meaningful morphological features associated with specific classes, two pathologists were asked to identify features that change during the transition to a counterfactual image. In these experiments, class labels for both the original and the counterfactual images were provided, and experts were asked to identify histopathological features that change during the transition. They were given 16 examples for each classifier (8 examples per class), each containing two top predictive tiles from the same patient who was predicted correctly by the model.

Furthermore, to formally quantify morphological changes for the MSI classifier, we conducted a structured survey with predefined colorectal features, mimicking a typical AI-assisted pathology workflow in which only highly attended regions are reviewed. For this, two expert pathologists independently reviewed paired original and counterfactual image tiles. For each transition, four tiles were shown in sequence (original → two intermediate diffusion steps → counterfactual), and pathologists were asked to indicate morphological features present in the leftmost (original) and rightmost (counterfactual) images. A predefined checklist of colorectal cancer histopathology features was provided (e.g., glandular architecture, necrosis, inflammatory infiltrate), and experts were instructed to select all applicable features for each image. The survey consisted of 52 image transitions (13 patients per class × 2 top-contributing tiles per patient), resulting in two feature annotations (original and counterfactual) per transition.

Finally, to additionally evaluate the realism of generated counterfactual images, a pathologist and a pathology trainee supervised by a board-certified pathologist reviewed a total of 300 images (150 real and 150 synthetic, randomly selected) in a blinded setting. For each task, 30 real tiles (15 per class) were randomly selected from the test set, and corresponding counterfactuals were generated by transitioning each tile to the opposite class. The pathologists were presented with a mixed set of tiles and asked to classify each one as either real or synthetic. The proportion of correctly identified images served as a measure of the diffusion model’s ability to generate realistic and visually convincing histopathology images.

Grad-CAMs were generated by backpropagating from the target class logit through the encoder’s last convolutional layer, computing gradient-weighted activations, and overlaying the resulting heatmap on the input tile.

### Independent Classifiers for Counterfactual Evaluation

To independently assess whether the MoPaDi-generated counterfactuals produce morphologically consistent class shifts, we employed independent classifiers trained on histopathology features extracted from large-scale foundation models. These classifiers were based on the STAMP (Solid Tumor Associative Modeling in Pathology) framework (41), which integrates pre-trained vision encoders with MIL for patient-level prediction. Using the same TCGA-CRC training split, we trained three classifiers with features extracted with UNI2 (42), Virchow2 (43), and Conch (44) models. For each encoder, features were aggregated at the patient level using the STAMP MIL transformer. All classifiers were trained to distinguish between MSIH and the absence of this biomarker (hereafter referred to as nonMSIH) using a categorical cross-entropy loss. Each classifier was applied to both the original and counterfactual images produced by MoPaDi. Predictions were obtained independently for every manipulation amplitude (0.02–0.08), yielding per-tile output for both classes. We quantified classifier agreement with the intended manipulation direction by analyzing classifier output trajectories across increasing manipulation amplitudes. Specifically, for each tile and encoder, we measured the change in predicted target-class probability (Δ*P*(*target*)) as a function of amplitude. Aggregated trajectories and mean Δ*P*(*target)* values were visualized to assess whether counterfactuals consistently increased the likelihood of the target phenotype across encoders.

### Quantitative Assessment of Cellular and Morphological Changes

Pretrained nuclei segmentation and cell type classification models from DeepCMorph (45) were applied to both original and counterfactual images to quantify cell type proportions and morphological characteristics. DeepCMorph models were pretrained on varying resolution H&E tiles (224 x 224 px) from diverse datasets to classify nuclei of six cell types: epithelial, connective tissue, lymphocytes, plasma cells, neutrophils, and eosinophils. Although originally optimized for cancer and tissue type classification, the models capture biologically meaningful cellular features that enable accurate segmentation-based quantification.

For epithelial cells, instance segmentation masks were derived by performing watershed-based splitting of touching objects in a semantic segmentation mask, followed by an additional binary erosion to split remaining touching nuclei, connected component labeling, and binary dilation to restore the original nucleus size. Nuclei smaller than 30 pixels were discarded as artifacts. Connective tissue nuclei, which are more elongated and spatially separated, were segmented directly from the predicted masks without further postprocessing.

From the resulting instance masks, we extracted morphological and intensity-based measurements, i.e., area, mean intensity, and eccentricity, using the *scikit-image* library. To capture spatial tissue organization, we computed the entropy of cellular spatial distributions by binning cell centroids into a 2D histogram, normalizing it to a probability distribution, and calculating Shannon entropy, which was further normalized by the maximum possible entropy given the histogram resolution. We also quantified cell counts per type and computed cell-type fractions (relative proportions).

For each patient, changes were assessed by comparing medians of these measurements between the counterfactual and original images. The direction of change was signed such that positive values indicate a shift toward MSIH morphology. Cohort-level summaries were computed as median signed deltas across patients, with interquartile ranges (IQRs) as variability estimates. Statistical significance was evaluated using two-sided Wilcoxon signed-rank tests.

To quantitatively compare changes in cell type proportions between original and counterfactual images of the opposite class (i.e., manipulated to look like the compared images’ class), we performed statistical testing at the patient level. For each patient and cell type, the mean proportion across analyzed tiles was computed separately for the original and manipulated images. Differences between original and counterfactual distributions were assessed using the two-sided Mann-Whitney U test, as the data were non-normally distributed and unpaired across patients. *P*-values were adjusted for multiple comparisons across cell types using the Bonferroni correction (family-wise *α* = 0.05). Effect sizes were quantified as the Hodges-Lehmann estimator of the median shift (in percentage points) with 95% percentile-bootstrap CI and complemented by Cliff’s delta (*δ*), interpreted using conventional thresholds (|δ| < 0.147 = negligible, < 0.33 = small, < 0.474 = medium, ≥ 0.474 = large).

### Data and Code Availability

Digital histopathology WSIs from TCGA, together with the clinical data used in this study, are available at https://portal.gdc.cancer.gov and https://cbioportal.org. NCT-CRC-HE-100K, CRC-VAL-HE-7K (32) is publicly available at https://zenodo.org/rec-ords/1214456. MoPaDi code and instructions to download trained models are publicly available on GitHub and Hugging Face at https://github.com/KatherLab/mopadi and https://huggingface.co/KatherLab/MoPaDi.

## Results

### Diffusion Autoencoders Faithfully Represent and Reconstruct Histopathology Images

MoPaDi is a diffusion-based model that preserves global tissue architecture while capturing fine cellular detail in its latent representation, enabling accurate reconstruction of histopathology images. To assess the fidelity of these latent representations, we first quantified reconstruction performance across multiple settings by training four independent models on the following datasets: TCGA-CRC, TCGA-BRCA, TCGA Pan-cancer, and the NCT-CRC-HE-100K dataset. We observed consistently high image reconstruction performance, reflected in low MSE and high MS-SSIM values across datasets (**Table 1**). Qualitative review further demonstrated near-identical reconstructions, with only subtle artifacts occasionally accentuated by the compression-decompression process (**Fig. S2**). In comparison to GAN-based methods, MoPaDi is able to achieve near-perfect reconstructions due to the encoded structural information of the original image in the latent space (**Fig. S3**).

**Table 1.** Evaluation of diffusion autoencoders trained on different datasets. The reconstruction quality of encoded images was evaluated using multi-scale structural similarity index measure (MS-SSIM) and mean square error (MSE), which perform pixel-by-pixel comparisons of 1,000 reconstructed images. The ability of the diffusion model’s latent space to capture meaningful information was further assessed by generating 10,000 synthetic histopathology images and computing the Fréchet Inception Distance (FID) score between the generated and held-out test set images to evaluate their quality.

| Model | MS-SSIM ↑ | MSE ↓ | FID score ↓ |
| --- | --- | --- | --- |
| NCT-CRC-HE-100K | $0.968 \pm 0.036$ | $(0.4 \pm 0.5) \cdot 10^{-3}$ | 36.95 |
| TCGA Pan-cancer | $0.992 \pm 0.001$ | $(0.8 \pm 0.2) \cdot 10^{-3}$ | 4.71 |
| TCGA-CRC | $0.967 \pm 0.060$ | $(8.1 \pm 14.0) \cdot 10^{-3}$ | 16.85 |
| TCGA-BRCA | $0.966 \pm 0.062$ | $(4.5 \pm 11.3) \cdot 10^{-3}$ | 17.85 |

We further assessed the encoder’s ability to capture meaningful histopathological features by training DDIMs on feature vectors extracted by the MoPaDi encoders for each dataset, enabling the sampling of synthetic feature vectors for downstream image generation. These models produced high-quality synthetic images, indicated by low FID between a held-out set of original images and 10,000 generated synthetic images (**Fig. S4**) for most datasets (**Table 1**). Values in the low tens generally reflect strong fidelity in histopathology. The highest FID score of 36.95 was obtained by the model trained on NCT-CRC-HE-100K when evaluated on the CRC-VAL-HE-7K dataset, reflecting the greater histologic heterogeneity of this dataset. Together, these results demonstrate that our diffusion autoencoders effectively capture key image features and reconstruct histopathology tiles, providing a robust foundation for downstream counter-factual image generation.

### Tissue Type Transformations via Latent Editing Exhibit Meaningful Patterns

After validating MoPaDi’s reconstruction performance and latent structure, we next examined whether the model can manipulate these representations to generate biologically plausible counterfactual images across tissue types. For this, we first trained a diffusion autoencoder alongside a feature extractor on the NCT-CRC-HE-100K dataset, representing nine tissue classes in colorectal cancer (**Fig. S5A**). Subsequently, we trained a logistic regression classifier on the extracted features to jointly predict all classes. For the generation of counterfactual explanations, we focused on the clinically most relevant pair: normal colon mucosa and dysplastic/tumor epithelium. However, counterfactual images can also be generated for other tissue types predicted by the classifier (representative examples for adipose, smooth muscle, mucus, debris, lymphocytes, and cancer-associated stroma counterfactuals are shown in **Supplementary Fig. S6**). Evaluated on the independent CRC-VAL-HE-7K dataset, the classifier reached an area under the curve (AUC) of 0.91 ± 0.01 for normal colon mucosa and 0.98 ± 0.01 for tumor epithelium (95% CI). AUC values for the remaining classes ranged from 0.79 to 1.00 (**Fig. S5B**). We used the classifier’s class-direction vector *v* to shift the latent features toward a selected target class *X*. The manipulated features then conditioned the diffusion-based decoder, enabling us to generate counterfactual images for tiles from the CRC-VAL-HE-7K dataset at increasingly larger manipulation amplitudes (*α*) (representative example of generated transition to a counterfactual image in **Fig. 2A**). When visualized in two-dimensional space, features extracted by MoPaDi’s encoder from the training dataset clustered by tissue type, suggesting that the model captured class-specific representations (**Fig. 2B**). Manipulated features also shifted closer to the opposite class in the lower-dimensional space (**Fig. 2B** zoomed-in plot), particularly when features are near the boundary between classes. To quantify the effectiveness of counterfactual generation, we computed the percentage of manipulated tiles that were predicted as the opposite class across increasing amplitudes (**Fig. 2C**). We observed that normal colon mucosa to tumor epithelium transitions generally required lower amplitudes to flip the classifier prediction, consistent with the lower AUC for the healthy class.

**Figure 2.**
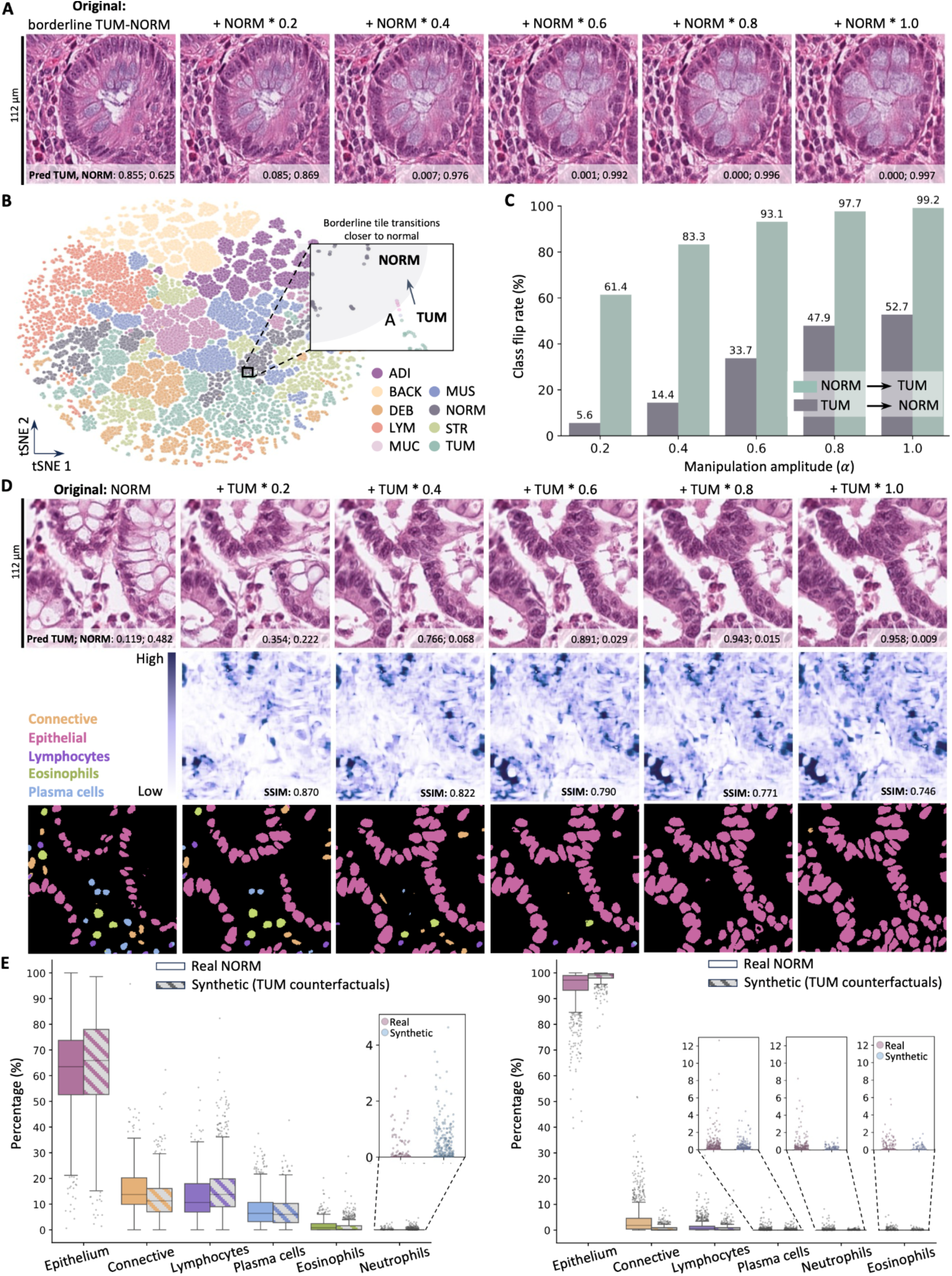
Results for counterfactual image generation for the CRC-VAL-HE-7K dataset. **(A)** Tissue-type classifier-guided counterfactuals of a dysplastic tile, generated by shifting its feature vector toward the predicted healthy colon mucosa class. Changes reflect what the model requires to flip the prediction, with increasing manipulation amplitude shown above each image. **(B)** t-SNE projection of the feature extractor’s latent space (training set), colored by class: adipose (ADI), background (BACK), debris (DEB), lymphocytes (LYM), mucus (MUC), smooth muscle (MUS), healthy colon mucosa (NORM), cancer-associated stroma (STR), colorectal adenocarcinoma epithelium/dysplastic tissue (TUM). The features of dysplastic tile in Panel **A**, initially near the boundary between TUM (aqua) and NORM (gray), shift toward the healthy cluster (pink points in the zoomed-in plot; gray area – arbitrary healthy mucosa zone) as manipulation amplitude increases. **(C)** Percentage of manipulated tiles predicted as the target class at increasing manipulation amplitudes (α). NORM to TUM manipulations require lower amplitudes to flip predictions compared to TUM to NORM, which is consistent with the classifier’s performance, as the classification performance is lower for NORM. **(D**) Counterfactuals of a healthy mucosa tile shifted toward dysplastic epithelium, rated highly realistic in a blinded user study. **Below**: pixel-level difference maps (darker regions indicate greater changes), showing the most change in gland regions, and Structural Similarity Index Measure (SSIM) values, which quantify the similarity between the original and manipulated tiles. **Bottom** row shows segmented and classified nuclei: pink (epithelial), orange (connective tissue), blue (plasma), deep purple (lymphocytes), aqua (neutrophils), green (eosinophils). **(E)** Tile-level differences in cell-type fractions between (**right)** real NORM tiles (*n* = 741) and synthetic (counterfactual images for TUM class, *n* = 1,233) (refer to **Table S3** for all statistical details). (**Left)** Tile-level differences in cell-type fractions between real TUM tiles (*n* = 1,233) and synthetic (counterfactual images for NORM class, *n* tiles = 741).

We conducted a blinded evaluation study with two pathologists to further assess the quality of counterfactual images. Each of them was shown 30 transitions to a counterfactual image (15 for tumor epithelium and 15 for normal colon mucosa) and asked to identify morphological changes. For example, counterfactual images showed that the tile was classified as tumor epithelium rather than normal colon mucosa due to the absence of healthy, well-ordered goblet cells, which appear in the counterfactual image (**Fig. 2A**). Conversely, the reverse counterfactual pair (normal colon mucosa to tumor epithelium) displayed loss of goblet cells and connective tissue, along with the appearance of hyperchromatic nuclei, revealing plausible morphological changes that would shift the classification toward colorectal cancer epithelium (**Fig. 2D**).

We further evaluated the quality of the generated images quantitatively by analyzing the distribution of six cell types in real and synthetic images (i.e., epithelial cells, connective tissue cells, lymphocytes, plasma cells, eosinophils, and neutrophils) (**Fig. 2E**). The results indicated that, when comparing real and synthetic images, cell-type composition differences were generally small or negligible (**Table S3**). Across the six celltype comparisons, all cell types, except plasma cells, showed significant tile-level differences between original healthy colon mucosa tiles (*n* = 741) and tumor tiles manipulated to be classified as healthy class (*n* = 1,233, *p* < 0.05, Mann-Whitney U, Bonferroni corrected), but all observed effect sizes were small (Hodges-Lehmann median shift ≤ 3 percentage points). When comparing real tumor epithelium tiles (*n* = 1,233) and their counterparts generated from healthy mucosa tiles (*n* = 741), the analysis revealed small but statistically significant shifts in connective tissue, lymphocytes, plasma cells, and epithelium fractions (*p* < 0.05), but with a large effect size only for the epithelial cells fraction. On average, the epithelial cell proportion was 1.57 percentage points higher in the synthetic images than in the real images (95% CI: –1.83 to – 1.34), and 1.27 percentage points lower for connective tissue cells (CI: 1.07–1.47), suggesting that MoPaDi has captured epithelial cell abundance as a key feature distinguishing malignant epithelium.

Collectively, these findings demonstrate that our model captured and could generate key histological features associated with normal colon mucosa and dysplastic epithelium, and that counterfactual images can serve as intuitive visual tools to reveal which morphological features influence the classifier’s predictions.

### Counterfactuals Efficiently Capture Microsatellite Instability-Related Morphology

Having established that MoPaDi generates biologically meaningful counterfactuals for tissue classification, we next evaluated its ability to predict a molecular biomarker from histopathology and to reveal the associated morphological patterns. We explored the link between genotype and phenotype by investigating counterfactual explanations for MSI in the TCGA-CRC cohort. A 5-fold cross-validation was performed with MIL to predict MSIH versus nonMSIH, achieving a mean AUC of 0.73 ± 0.08 and a mean average precision (AP) of 0.40 ± 0.09, with lower AP reflecting high-class imbalance. We then generated counterfactual transitions to expose the morphology captured by the classifier. Pathologists identified that the tiles contributing most strongly to the model’s predictions for MSIH cases typically contained inflammatory cells, atypical epithelium lacking normal bowel crypt structure, debris, high-grade nuclear pleomorphism, and, in some cases, medullary or mucinous morphology and signet-ring cells. Counterfactuals were generated on the top-5 contributing tiles per WSI. Counterfactual transitions indicated that an MSIH tile would be predicted as nonMSIH if it displayed more organized glandular architecture, reduced mucin content, and a less solid, sheet-like growth pattern when the original tile displayed a medullary growth (**Fig. 3A, Fig. S8**). Conversely, counterfactual edits indicated that nonMSIH tiles would be classified as MSIH if they showed reduced glandular architecture, more solid growth patterns, increased mucinous appearance, and a more vacuolated appearance (**Fig. 3B, Fig. S9**). Pathologists’ survey results are summarized quantitatively in **Fig. 3C**. Furthermore, many of the generated images for both classes revealed center-specific batch effects through visible color changes. Quantitative measurements supported these observations, showing significant changes in mean nuclear intensity for both epithelial and connective tissue cells (**Fig. S10C**). These findings motivated us to further analyze the relative contributions of style and morphology to model decisions (see the following section).

**Figure 3.**
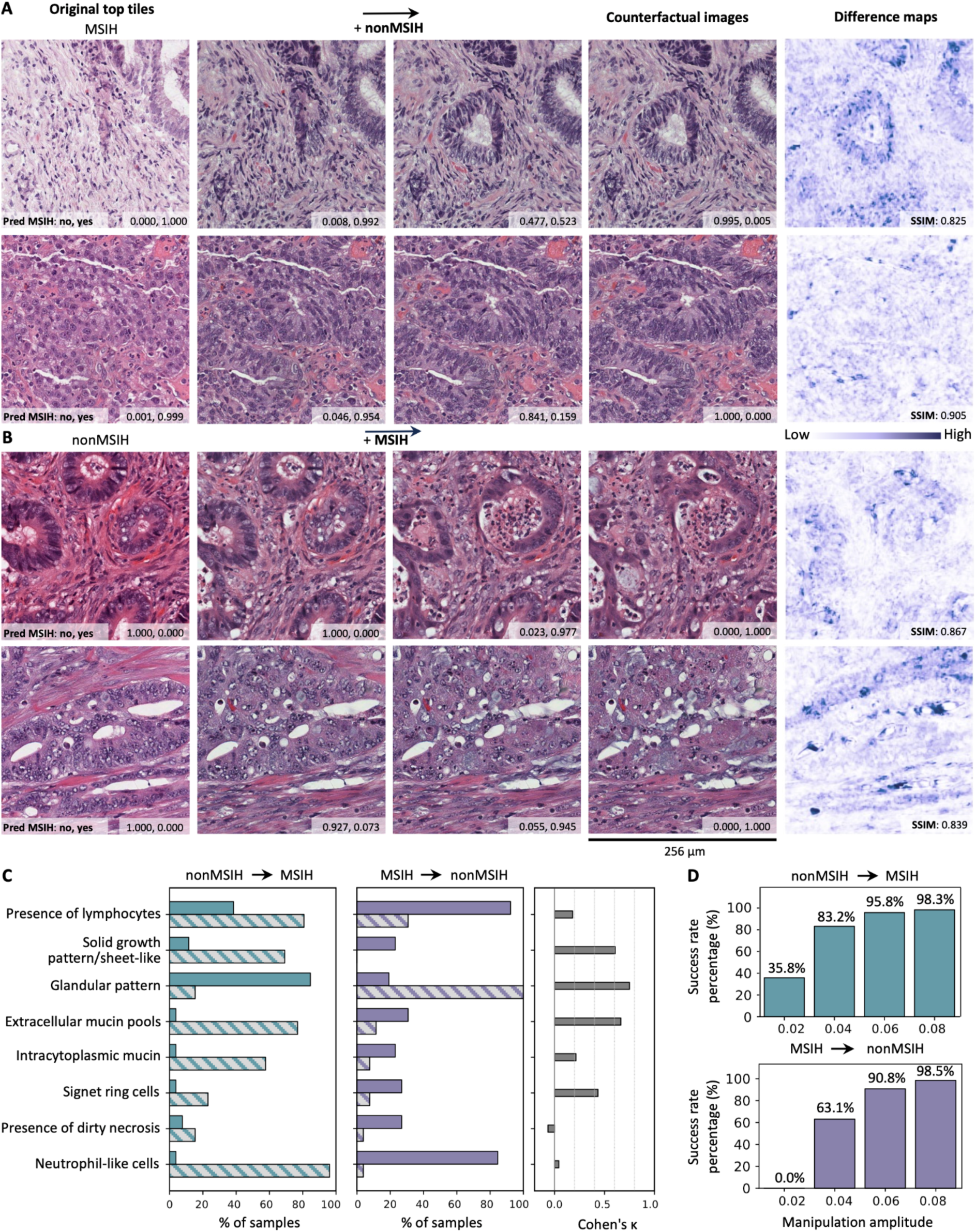
Interpretation of the molecular biomarker (microsatellite instability [MSI]) classifier in colorectal cancer through counterfactual image analysis. **(A)** Representative examples of MSI-high (MSIH) tiles transitioning to their counterfactual nonMSIH image. Pixel-wise difference images compare the original image with the most manipulated one. **(B)** Representative examples of nonMSIH tiles transitioning to their counterfactual MSIH image. Pixel-based difference maps highlight changes in nuclei, stroma, cell density, and tissue architecture. **(C)** Results of the pathologist survey. Two pathologists were shown 52 transitions from an original image to a counterfactual one (13 MSIH patients and 13 nonMSIH patients, 2 top contributing tiles from each patient) and asked to identify morphological features that were present on different sides. **(D)** Quantitative evaluation of counterfactual image generation effectiveness. Bars show the percentage of successful manipulations (i.e., predicted class equals target class) across different manipulation amplitudes. SSIM – structural similarity index measure.

We further assessed changes in cell-type composition and cell-level morphological metrics, including nuclear area, solidity, or eccentricity, during counterfactual transitions (**Fig. S10**). Counterfactual edits from MSIH to nonMSIH exhibited a reduction in immune cells and a corresponding increase in epithelial and connective tissue fractions, whereas nonMSIH to MSIH transitions displayed the opposite trend, with a significant increase in immune cells. These shifts align with the well-established immune-rich microenvironment of MSIH tumors and reinforce that MoPaDi captures biologically meaningful tissue-level features. Notably, when comparing cell-type fractions between real images of a given class and counterfactual images manipulated toward that class, no significant differences were observed, suggesting that MoPaDi accurately captured tissue composition and generated class-consistent representations.

Additionally, we assessed both the robustness and effectiveness of our counterfactual generation approach across increasing manipulation amplitudes (*α*). First, we observed that most counterfactuals successfully reached the target class even at low *α* values, demonstrating efficient movement along the model’s decision boundary (**Fig. 3D**). Second, to assess robustness, we examined whether independent histopathology classifiers agree with the direction of the counterfactual changes. We applied three state-of-the-art foundation model-based classifiers (UNI2, Conch, and Virchow2) to original and counterfactual tiles and quantified changes in MSI prediction confidence across increasing manipulation amplitudes. Consistent shifts in MSI probability in the intended direction across all classifiers confirmed that counterfactual trajectories are supported by external models and not specific to a single classifier (**Fig. 4A, Fig. S11**).

**Figure 4.**
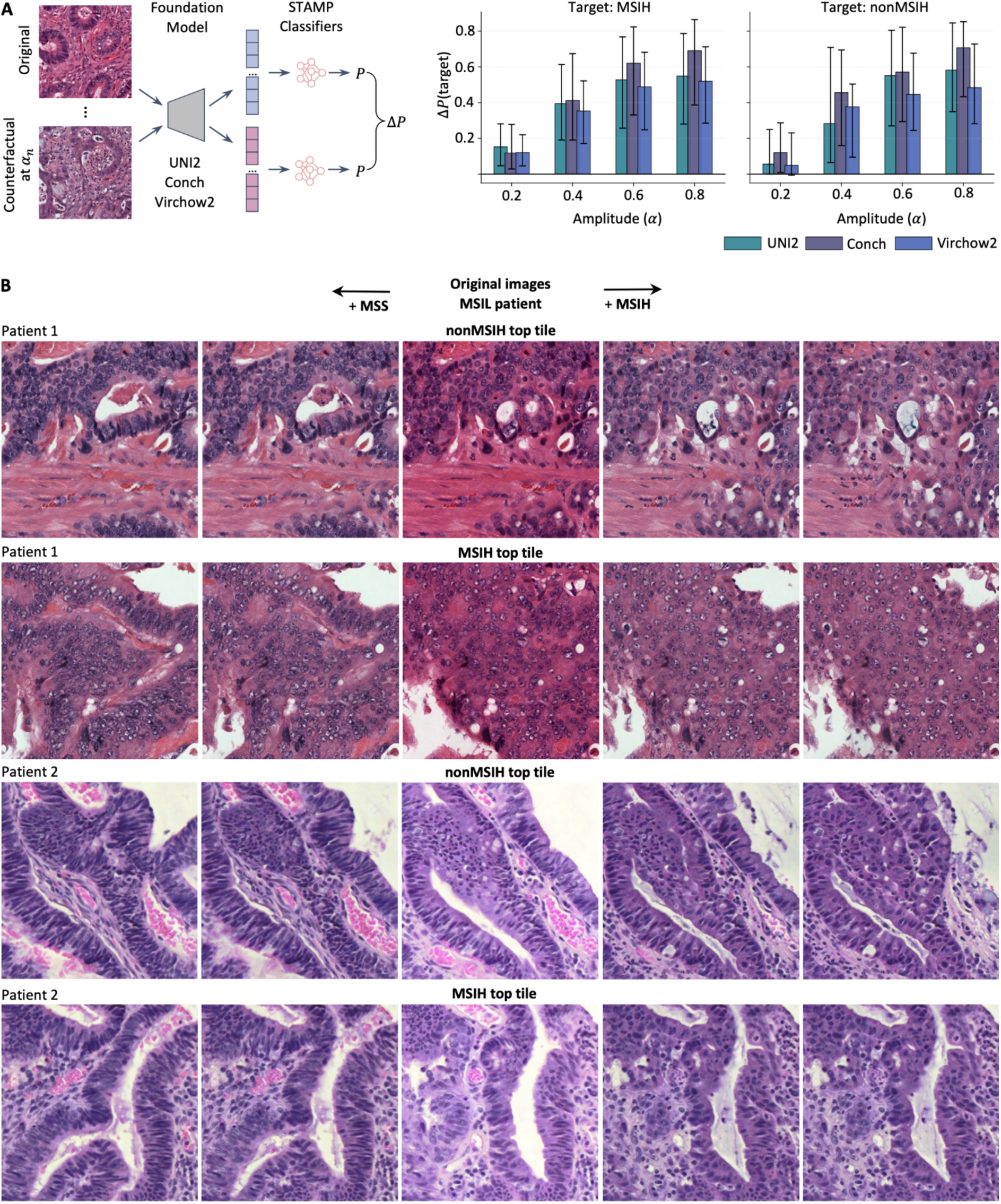
Independent classifiers validation and representative bidirectional counterfactual transformations in a multiclass setting. **(A)** External classifier responses to counterfactual morphing. Counterfactual images were generated at increasing morphing amplitudes (*α*) with MoPaDi, then encoded with three foundation models (UNI2, Conch, Virchow2). Independent classifiers were trained on the corresponding encoders’ features extracted from all TCGA-CRC and then used to predict the target probability *P(target)* on both the original and counterfactual tiles. The resulting change Δ*P(target)* reflects how strongly the morphing affected class evidence. Bars show the median Δ*P(target)* with percentile-based variability across tiles. **(B)** Representative examples of bidirectional counterfactual explanations for microsatellite instability-low (MSIL) patients. We defined MSIL by fitting a two-component Gaussian mixture model to the log-transformed distribution of total MSI events and using the intersection of the two components as the cutoff separating MSIL from MSIH samples.

Generated images should not only flip the class but also remain biologically plausible. We evaluated perceptual realism through a blinded review by two pathologists, each assessing 300 tiles (150 real, 150 synthetic) as either real or synthetic (**Fig. S12**). Across all datasets, 26.7%–63.3% of synthetic tiles were misclassified as real, demonstrating that MoPaDi produced images that often appear realistic to experts.

We next evaluated whether our framework generalizes to the more nuanced three-class MSI setting. MSI status is clinically categorized as MSS, MSIL, or MSIH, based on the fraction of tested microsatellite loci in which instability is detected. To qualitatively assess MoPaDi in this multiclass context, we generated counterfactual images for representative MSIL patients, selecting tiles with high model attention toward either the nonMSIH or MSIH class (**Fig. 4B**). Consistent with previously described results, generated edits towards MSIH showed progressive emergence of features associated with microsatellite instability, including disappearance of glandular architecture (e.g., more solid growth pattern), and more intratumoral lymphocytes, while the MSS direction restored gland formation and reduced mucin content. These bidirectional transitions demonstrate that MoPaDi modifies tissue morphology in a class-specific and interpretable manner, confirming that the model has captured histologically relevant representations of MSI status.

These findings, from pathologists’ observations of architectural changes in counter-factual transitions to quantitative analyses of cellular composition, reveal candidate morphological cues the classifier uses to distinguish MSIH tumors from nonMSIH, enhancing the interpretability of the prediction process.

### Style-Morphology Analysis Disentangles Staining from Structural Effects

To disentangle which visual aspects of the tissue drive the model’s predictions, we performed a style-morphology decomposition of real and counterfactual image pairs. This analysis aimed to separate the overall prediction change (Δ*f*) into contributions from staining style and tissue morphology, thereby quantifying their relative influence on model decision-making (**Fig. 5**).

**Figure 5.**
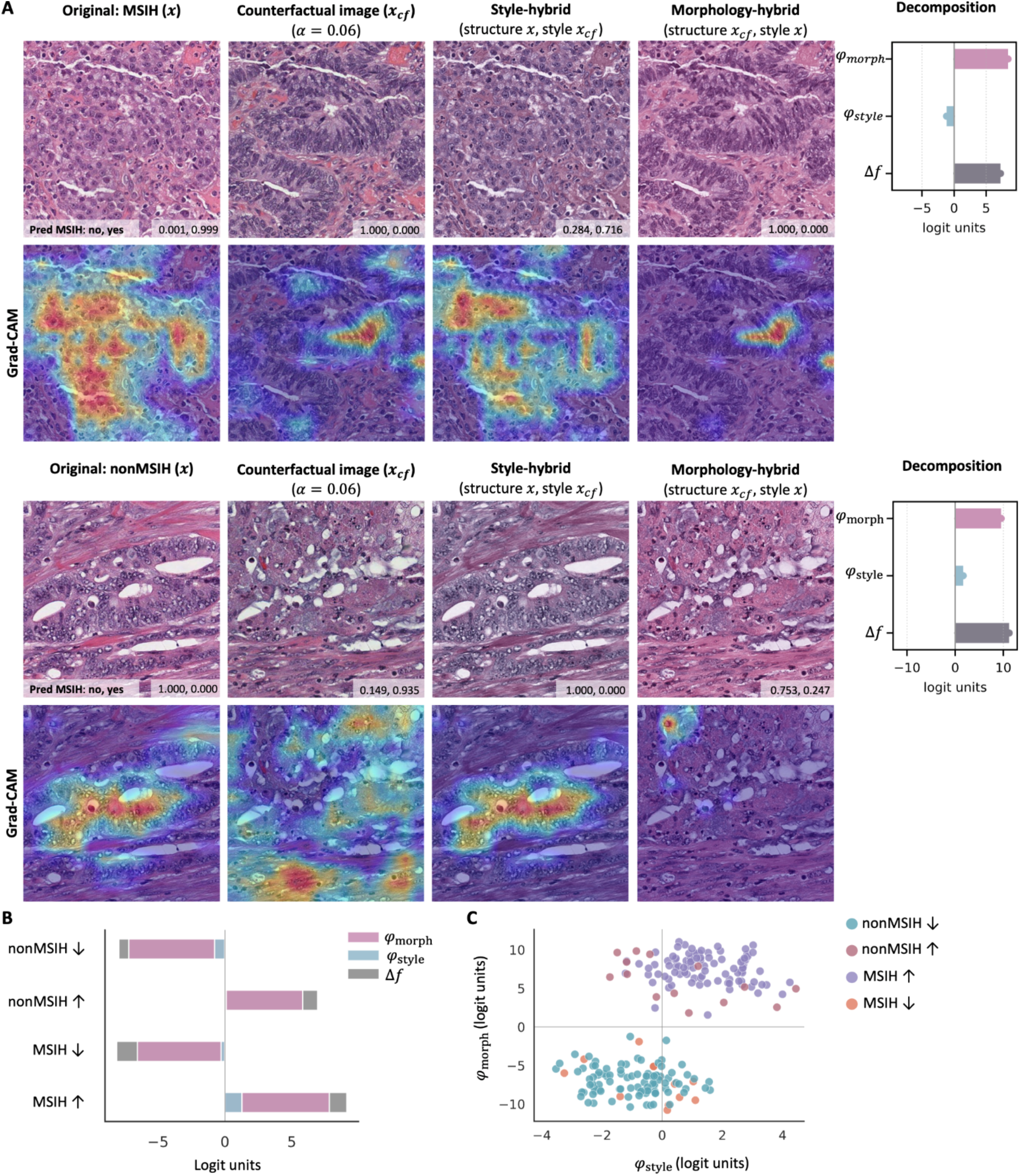
Style-Morphology decomposition for disentangling structural and staining effects on microsatellite instability (MSI) prediction changes. (**A**) Representative examples of counterfactual manipulation between MSI-high (MSIH) and nonMSIH classes. For each original image *x* and its counterfactual *x_cf_*, style-hybrid (*x_style_*) and morphology-hybrid (*x_morph_*) images were generated using Vahadane stain transfer. Each hybrid isolates the effect of either stain or morphology while controlling for the other. The right-hand bars show the Shapley-style decomposition of the logit change (Δ*f*) into stain (*φ_style_*) and morphology (*φ_morph_*) contributions, demonstrating that morphological differences dominate the model’s predictions. Gradient-weighted Class Activation Mapping (Grad-CAM) visualizations below highlight coarse, class-correlated regions but offer limited feature specificity under MIL. In contrast, MoPaDi produces class-directed ‘what-if’ edits that reveal which morphology must change for the logit to flip, and what is the influence of style. **(B)** Decomposition results across test-set patients, showing median contributions of *φ_style_*, *φ_morph_,* and total (Δ*f*) for manipulations toward (↑) and away from (↓) each class. **(C)** Scatter plot of morphology versus style contributions per patient, illustrating consistent dominance of morphological effects across both manipulation directions and classes.

We aggregated the decomposition results over all counterfactual tiles to quantify the overall contribution of style and morphology to the MSIH logit, as class transitions often include changes in staining. Across all colorectal cancer test-set patients (*N* = 108), manipulations toward the MSIH class yielded a median logit increase (Δ*f*) of +8.9 (IQR 7.2–10.1). The morphology-related component (*φ_morph_*) accounted for most of this shift (median +7.4, IQR 5.7–8.7), while the stain-related component (*φ_style_*) was comparatively small (+1.1, IQR 0.4–2.4). Morphological features dominated in all patients (morphology-dominant fraction reached 1.0), and the two components were generally reinforcing (conflict fraction was close to 0). These findings indicate that model decisions for MSIH status are primarily driven by morphological cues rather than staining variation.

We next analyzed counterfactual manipulations toward the nonMSIH class, aggregating decomposition results across all test-set patients. This analysis quantifies how the model’s MSIH evidence decreases when images are driven toward nonMSIH-like appearances. The manipulation resulted in a median logit change (Δ*f*) of -7.6 (IQR -9.0 to -6.3), reflecting a consistent reduction of MSIH evidence. Again, the morphology-related component accounted for the majority of this decrease (median -6.6, IQR -8.2 to -5.0), whereas the stain-related component contributed only modestly (median -0.7, IQR -1.9 to -0.1). Morphological effects dominated in all patients (morphology-dominant fraction = 1.0), with a small subset (conflict fraction of 0.2) showing opposing effects between style and morphology.

To further investigate whether the decomposition patterns depend on the ground-truth MSIH status of each sample, we stratified patients into MSIH and nonMSIH groups and repeated the aggregation separately. Across 109 colorectal cancer patients, bidirectional decomposition analysis revealed consistent morphological dominance in both MSIH and nonMSIH transitions. Manipulations toward MSIH increased model confidence (Δ*f =* +9.1 [IQR 7.8–10.5]), primarily through morphology-related components *φ*_morph_ *=* +7.8 [6.3–8.9] with minor staining influence (*φ*_style_ = +1.3 [0.6–2.6]). Conversely, transitions away from MSIH reduced confidence (Δ*f* = -8.1 [-10.0 to -6.3]), again morphology-driven (*φ*_morph_ *=* -6.5 [-9.1 to -5.1]). For true nonMSIH samples, manipulations toward nonMSIH and away from nonMSIH showed symmetric logit shifts (+6.9 [5.4–9.0] and –7.9 [-9.2 to -6.7], respectively), both dominated by morphological effects (*φ*_morph_ *=* 5.8 [4.0–8.3] and -7.2 [-8.3 to -6.1], respectively). Morphology dominated in nearly all patients (fraction ≈ 1.0), and style–morphology conflicts were infrequent (0.0–0.3).

Together, these results show that morphological structure is the primary driver of MSIH discrimination, with staining variations exerting a minor and sometimes opposing influence.

### Pan-cancer Evaluation Reveals Morphological Changes Across Cancer Types

We next evaluated MoPaDi across multiple cancer types to assess whether the framework generalizes beyond colorectal cancer and reveals biologically plausible morphology changes in diverse tumor contexts. We started by training a linear MoPaDi classifier to distinguish two hepatobiliary cancer types, hepatocellular carcinoma and cholangiocarcinoma, and generating related counterfactuals to determine what features the classifier captured. A logistic regression model trained on features extracted from hepatocellular carcinoma and cholangiocarcinoma cases in the TCGA Pan-cancer dataset achieved an AUC of 0.90 ± 0.01 (95% CI) and an AP of 0.57 ± 0.03 on the heldout test set. The lower AP is attributed to the imbalanced dataset, with only 10% of patients having cholangiocarcinoma (refer to **Table S2** for patient numbers). Averaging tile-level predictions to patient-level yielded an AUC of 0.94 ± 0.10, indicating an effective distinction between hepatobiliary cancer types. A qualitative analysis of the generated counterfactual images revealed that the model accurately captured representative morphological features for both classes. Hepatocellular carcinoma tiles manipulated to resemble cholangiocarcinoma displayed a more glandular pattern, reduced eosinophilic cytoplasm, and lesser trabecular architecture (**Fig. 6A**). Meanwhile, cholangiocarcinoma tiles manipulated to resemble hepatocellular carcinoma displayed a more trabecular growth pattern with loss of glandular structure, eosinophilic cytoplasmic inclusions, possible Mallory bodies, and reduced stromal tissue (**Fig. 6B**). These distinct morphological changes in counterfactual images demonstrate the model’s ability to capture hepatobiliary cancer characteristics while enhancing the classifier’s interpretability.

**Figure 6.**
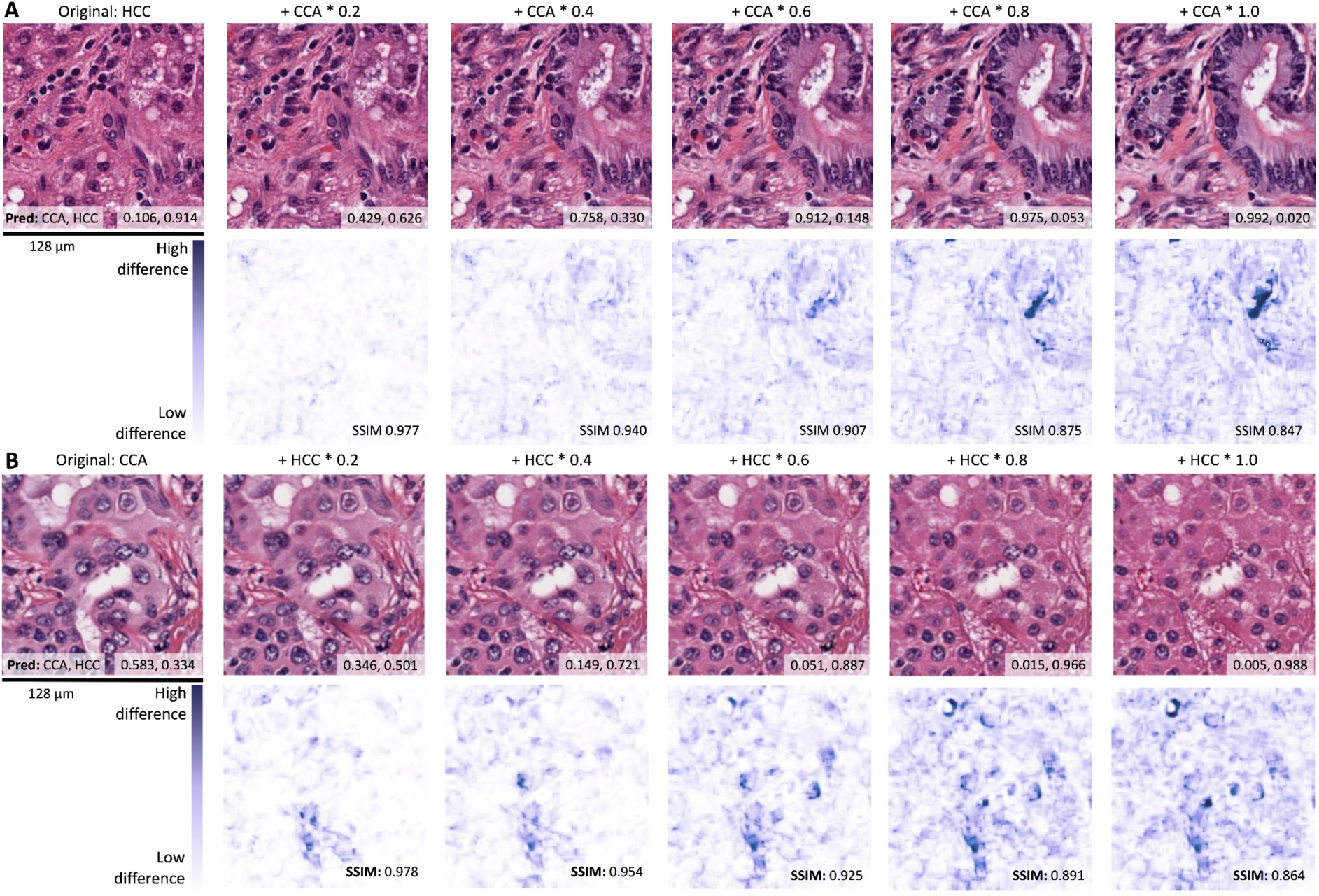
Counterfactual image examples generated from the liver cancer type classifier. (**A**) Representative example of hepatocellular carcinoma (HCC) tile transitioning to its counterfactual cholangiocarcinoma (CCA) image generated with a linear approach. Comparison to the MIL approach can be found in the supplementary material (**Fig. S13**). Difference maps display pixel-wise differences between the original and the synthetic tile. (**B**) Representative example of CCA tile transitioning to its counterfactual HCC image. SSIM, structural similarity index measure.

Given the complexity of weakly supervised WSI tasks, where histological phenotypes are visible only in select regions, we assessed classification with attention-based MIL of different cancer types: lung cancer (lung adenocarcinoma vs. lung squamous cell carcinoma), breast cancer (invasive lobular carcinoma [ILC] vs. invasive ductal carcinoma [IDC]), and the previously discussed hepatocellular carcinoma vs. cholangiocarcinoma. For the latter task, integrating MIL led to predictive performance marginally increasing compared to the tile-based linear approach, achieving a mean AUC of 0.96 ± 0.01 and an AP of 0.82 ± 0.07 in the 5-fold cross-validation. Top tiles from correctly classified patients displayed distinctive features: original hepatocellular carcinoma tiles showed cells arranged in sheets or trabecular patterns with lipid droplets, lacking glandular structures and fibrous stroma. In contrast, original cholangiocarcinoma tiles exhibited glandular structures and more fibrous stroma (**Fig. S13A**). Increasing manipulation amplitude (*α*) resulted in higher opposite-class predictions, with 100% of generated images being predicted as the opposite class at *α* = 0.08 (**Fig. S13B**). Counterfactual images guided by the MIL classifier resembled those generated using the linear approach with tile-level logistic regression (**Fig. S13C**). The cell-type composition did not differ significantly between counterfactual and real images, indicating that counterfactual edits preserved underlying tissue composition (**Fig. S13D**).

We further validated counterfactual image generation guided by the MIL classifier by training a lung adenocarcinoma vs. lung squamous cell carcinoma classifier, achieving a mean AUC of 0.91 ± 0.01 (with an improvement of 0.10 over the linear method) and an AP of 0.89 ± 0.02. Pathologists analyzed lung squamous cell carcinoma to lung adenocarcinoma counterfactual transitions (**Fig. 7A**). Increased glandular differentiation was noted in 4/8 and 5/8 patients by the two reviewers, respectively, and both observed reduced nuclear pleomorphism in 6/8 cases. Additional changes included increased vacuolization, clearer cytoplasm (each 4/8), and reduced tumor solidity (3/8). These observations suggest that lung adenocarcinoma-associated features emerge during counterfactual editing, and their absence contributes to the classification of tumors as lung squamous cell carcinoma. However, some counterfactual tiles retained lung squamous cell carcinoma features atypical for lung adenocarcinoma, such as prominent dyskaryotic changes, nuclear molding, and residual squamous differentiation (**Fig. S14A**), indicating that the model may prioritize other features when making classification decisions. Additionally, lung adenocarcinoma to lung squamous cell carcinoma transitions showed decreased glandular architecture, increased solid growth patterns or tumor cell nests (4/8; 5/8), and increased cell density (2/8; 5/8). Other features included lower atypia (1/8; 2/8), intercellular bridges, and a keratin pearl (**Fig. S14B**). Counterfactual images maintained largely stable cell type compositions across most cell types, with minor differences observed, notably in immune cells (**Fig. S14C**). Counterfactual image generation effectiveness remained high across both classes, with a total of 98.8% of counterfactual images at amplitude α = 0.06 being predicted as the opposite class relative to the original (**Fig. S14D**). Taken together, these data demonstrate that MIL classifier-guided counterfactual image generation works effectively not only for colorectal cancer.

**Figure 7.**
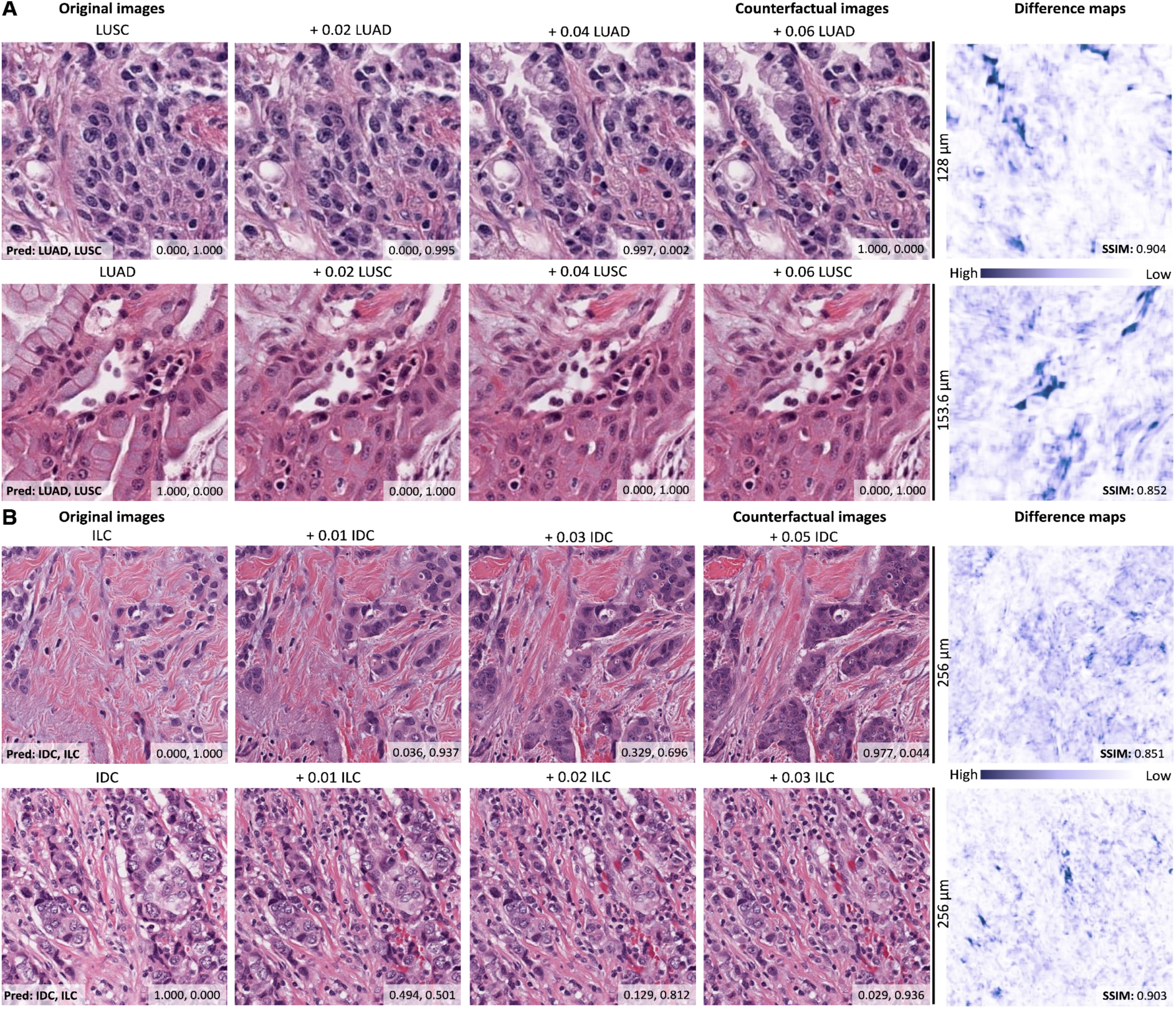
Counterfactual image examples generated from the breast and lung cancer type classifiers. (**A**) Representative examples of lung squamous cell carcinoma (LUSC) tile transitioning to its counterfactual lung adenocarcinoma (LUAD) image and vice versa. (**B**) Representative examples of invasive lobular carcinoma (ILC) tile transitioning to its counterfactual invasive ductal carcinoma (IDC) image and vice versa. Difference maps display pixel-wise differences between the original and the synthetic tile. SSIM, structural similarity index measure.

Finally, we investigated whether meaningful morphological features could be identified in counterfactuals for the classification of IDC and ILC in the TCGA-BRCA cohort. The trained classifier reached a mean AUC of 0.86 ± 0.02 and AP of 0.64 ± 0.04. In ILC to IDC counterfactual transitions (**Fig. 7B top**), the most consistently observed changes were increased hyperchromasia (5/8) and atypia (4/8), showing that the tiles were classified as ILC due to the absence of these features. Other notable features included a shift towards less single-cell arrangement without clear glandular pattern formation (2-3/8), larger tumor cells (3/8), and tumor cell arrangement in larger nests (2/8). Some top tiles contained only muscle with single tumor cells, making feature assessment not feasible, but potentially representing the diffusely infiltrative nature of these neoplasms. Meanwhile, IDC to ILC counterfactual transitions (**Fig. 7B bottom**) lacked glandular architecture (7/8), exhibited less prominent nucleoli (8/8), and exhibited a single-cell arrangement of tumor cells (8/8), suggesting that the classifier distinguishes IDC based on the prominence of nucleoli and growth patterns. Although generated images appeared as realistic ILC representations with 97.4% of counterfactuals predicted as the flipped class (**Fig. S15C**), the top contributing tiles for which counterfactuals were generated often lacked characteristic IDC features. However, despite the classifier’s moderate performance and inconsistent features in top tiles, we identified distinct morphological characteristics of ILC and IDC, indicating that the classifier captured meaningful distinctions between these cancer types.

## Discussion

Throughout this study, we developed a diffusion model – MoPaDi – capable of generating meaningful counterfactual images and enhanced it with MIL to handle WSIs. MoPaDi demonstrates how diffusion models can transform computational pathology from a passive descriptive tool into an instrument for virtual experimentation. Unlike traditional DL approaches that simply map images to predictions, MoPaDi enables pathologists to ask and answer “what if” questions about biological specimens through counterfactual image generation. While not implying biological causality, these counterfactuals provide model-internal simulations of morphological change.

The effectiveness of this approach was extensively validated across multiple experimental scenarios, spanning multiple tumor types and providing quantitative and qualitative evidence for the plausibility of generated images by using several orthogonal metrics. Our diffusion autoencoder achieved excellent image reconstruction quality (MS-SSIM) and generated realistic counterfactual images that were often indistinguishable from real histological samples in blinded pathologist evaluations. Furthermore, we provide evidence that MoPaDi captures and recapitulates key biological processes. For example, the overall trend for cell type distributions in generated images of colorectal cancer aligns with established cancer and adjacent normal tissue microenvironment characteristics (46). Tumor tiles showed high epithelial cell concentrations, especially in normal to tumor transitions, suggesting the model captures epithelial abundance as a key tumor characteristic, consistent with tumor biology (47). Similarly, for distinguishing liver tumor types and lung tumor types, our model recapitulated key biological properties, such as trabecular to glandular transitions and the amount of fibrous stroma in liver samples (48–50). Finally, we extended our analysis to molecular biomarkers by investigating MSI status in colorectal cancer. While achieving moderate performance compared to state-of-the-art models (3), MoPaDi revealed key morphological patterns associated with MSI status through counterfactual explanations. Even with a limited set of top tiles, which mimics a typical workflow in computational pathology studies, pathologists identified characteristic features such as signet-ring cells, mucinous, medullary patterns, inflammatory infiltration, and poor differentiation that are known to be associated with MSI in colorectal cancer (51–55). These findings show that DL models can be used not only for image-based classification tasks but also for controlled, biologically interpretable image manipulations. Unlike standard attention heatmaps typically used in computational histopathology, which highlight broad regions of interest and can lead to confirmation bias (6), our approach identifies specific features the model uses for predictions.

As another application of MoPaDi, we demonstrate its ability to disentangle morphological structure from staining style, revealing which visual aspects predominantly drive model predictions. Staining variations are a common source of bias in computational pathology, as DL models may rely on site- or scanner-specific color cues instead of biologically relevant morphology (56–58). By decomposing prediction changes into style- and morphology-related components, MoPaDi quantitatively distinguishes between these influences for each image. This analysis revealed that model decisions were morphology-driven, with staining exerting only a minor or opposing effect. Beyond improving interpretability, this disentanglement enables the identification of potential staining biases at a per-image level, providing a systematic approach to evaluate style-related confounders in digital pathology models.

Finally, to the best of our knowledge, MoPaDi is the first generative model that works in a MIL framework to generate counterfactual explanations for biomarkers that are only defined on the level of slide annotations. This capability is important because many important molecular and clinical features in computational pathology are only available at the whole-slide level. By combining diffusion models with MIL, we enable the interpretation of weakly-supervised learning problems that are common in clinical practice.

Our approach has several important limitations. Results are currently limited to tile-level analysis and require careful validation by domain experts to ensure biological plausibility. Each dataset requires training of a custom diffusion autoencoder model, which is computationally intensive and time-consuming, ranging from several days to weeks, depending on dataset complexity and hardware. Although the model typically generates realistic images, it sometimes fails to capture fine biological nuances. For instance, in colorectal cancer transitions, we observed cases where generated images lacked proper basement membrane structure or showed insufficient nuclear hyperchromatism. Additionally, while our cell type distributions generally align with known tumor biology, the model shows consistent biases in generating certain cell types over others. Future work could address these limitations by integrating pathology foundation models with diffusion-based counterfactual generation to improve both predictive performance and biological accuracy while maintaining the advantages of high-quality image synthesis and selective feature manipulation.

Looking forward, this work opens new possibilities for computational pathology. By enabling virtual experimentation, tools like MoPaDi could help researchers pre-screen hypotheses before conducting expensive and time-consuming wet lab experiments. This could be particularly valuable given increasing restrictions on animal testing and the challenge of translating animal model results to humans. Additionally, these methods could provide new insights into morphological features that correlate with outcomes, potentially revealing previously unknown biological mechanisms. As these methods continue to evolve, they may become an integral part of the cancer research toolkit, complementing traditional experimental approaches and accelerating the pace of discovery in cancer biology.

## Abbreviations

AI: Artificial Intelligence
AP: Average precision
AUC: Area under the curve
CI: Confidence intervals
DDIM: Denoising diffusion implicit model
DDPM: Denoising diffusion probabilistic model
DL: Deep learning
FID: Fréchet inception distance
GAN: Generative adversarial network
H&E: Hematoxylin and eosin
IDC: Invasive ductal carcinoma
ILC: Invasive lobular carcinoma
IQR: Interquartile range
MIL: Multiple instance learning
MPP: Micrometers per pixel
MS-SSIM: Multi-scale structural similarity index measure
MSE: Mean square error
MSI: Microsatellite instability
MSIH: Microsatellite instability-high
MSIL: Microsatellite instability-low
MSS: Microsatellite stable
ROC: Receiver operating characteristic
TCGA: The Cancer Genome Atlas
TCGA-BRCA: TCGA Breast Invasive Carcinoma
TCGA-CRC: TCGA Colorectal Carcinoma
WSI: Whole slide image

## Ethics Statement

This study was carried out in accordance with the Declaration of Helsinki. We retrospectively analyzed anonymized patient samples using publicly available data from “The Cancer Genome Atlas” (TCGA, https://portal.gdc.cancer.gov) and Zenodo. The Ethics Commission of the Medical Faculty of the Dresden University of Technology provided guidelines for study procedures.

### Acknowledgments

We gratefully acknowledge the GWK’s support for funding this project by providing computing time through the Center for Information Services and HPC (ZIH) at TU Dresden. We also thank the Gauss Centre for Supercomputing e.V. (www.gauss-centre.eu) for funding this project by providing computing time through the John von Neumann Institute for Computing (NIC) on the GCS Supercomputer JUWELS at Jülich Supercomputing Centre (JSC). Furthermore, we acknowledge the TCGA Research Network (https://www.cancer.gov/tcga), which generated the data on which the results shown in this manuscript are partly based. In accordance with the COPE (Committee on Publication Ethics) position statement of 13 February 2023, we hereby disclose the use of the following artificial intelligence models during the writing of this article. We used GPT-5 (OpenAI) to check spelling and grammar; all content was reviewed and verified by the authors.

## Author Contributions

LZ, JNK, and DT conceptualized the study. LZ and TL performed the experiments and developed the software. LZ, NGR, SF, and KJH performed results analysis. LZ prepared the original draft and created the figures. JNK acquired funding and supervised the study. All authors provided scientific input, reviewed, and edited the manuscript.

## References

1. Kather JN, Krisam J, Charoentong P, Luedde T, Herpel E, Weis C-A, et al. Predicting survival from colorectal cancer histology slides using deep learning: A retrospective multicenter study. PLoS Med. 2019;16:e1002730.

2. Zeng Q, Klein C, Caruso S, Maille P, Allende DS, Mínguez B, et al. Artificial intelligence-based pathology as a biomarker of sensitivity to atezolizumab-bevacizumab in patients with hepatocellular carcinoma: a multicentre retrospective study. Lancet Oncol. 2023;24:1411–22.

3. Wagner SJ, Reisenbüchler D, West NP, Niehues JM, Zhu J, Foersch S, et al. Transformer-based biomarker prediction from colorectal cancer histology: A large-scale multicentric study. Cancer Cell. 2023;41:1650–61.e4.

4. Calderaro J, Žigutytė L, Truhn D, Jaffe A, Kather JN. Artificial intelligence in liver cancer - new tools for research and patient management. Nat Rev Gastroenterol Hepatol. Springer Science and Business Media LLC; 2024;21:585–99.

5. Unger M, Kather JN. Deep learning in cancer genomics and histopathology. Genome Med. 2024;16:44.

6. Ghassemi M, Oakden-Rayner L, Beam AL. The false hope of current approaches to explainable artificial intelligence in health care. Lancet Digit Health. Elsevier BV; 2021;3:e745–50.

7. Huff DT, Weisman AJ, Jeraj R. Interpretation and visualization techniques for deep learning models in medical imaging. Phys Med Biol. 2021;66:04TR01.

8. Kim B, Wattenberg M, Gilmer J, Cai C, Wexler J, Viegas F, et al. Interpretability Beyond Feature Attribution: Quantitative Testing with Concept Activation Vectors (TCAV). In: Dy J, Krause A, editors. Proceedings of the 35th International Conference on Machine Learning. PMLR; 10-15 Jul 2018. page 2668–77.

9. Graziani, M., Lompech, T., Müller, H., & Andrearczyk, V. Evaluation and comparison of CNN visual explanations for histopathology. Proceedings of the AAAI Conference on Artificial Intelligence Workshops (XAI-AAAI-21), Virtual Event. page 8–9.

10. Amorim JP, Abreu PH, Fernandez A, Reyes M, Santos J, Abreu MH. Interpreting Deep Machine Learning Models: An Easy Guide for Oncologists. IEEE Rev Biomed Eng. 2023;16:192–207.

11. Guidotti R. Counterfactual explanations and how to find them: literature review and benchmarking. Data Min Knowl Discov. 2022; Available from: 10.1007/s10618-022-00831-6

12. Goyal Y, Wu Z, Ernst J, Batra D, Parikh D, Lee S. Counterfactual Visual Explanations. Proceedings of Machine Learning Research. 2019. page 2376–84.

13. Zhao W, Oyama S, Kurihara M. Generating Natural Counterfactual Visual Explanations. Proceedings of the Twenty-Ninth International Joint Conference on Artificial Intelligence. California: International Joint Conferences on Artificial Intelligence Organization; 2020. Available from: 10.24963/ijcai.2020/742

14. Han T, Žigutytė L, Huck L, Huppertz MS, Siepmann R, Gandelsman Y, et al. Reconstruction of patient-specific confounders in AI-based radiologic image interpretation using generative pretraining. Cell Reports Medicine. Elsevier; 2024 [cited 2024 Sep 7];5. Available from: https://www.cell.com/cell-reports-medicine/fulltext/S2666-3791(24)00434-8

15. DeGrave AJ, Cai ZR, Janizek JD, Daneshjou R, Lee S-I. Auditing the inference processes of medical-image classifiers by leveraging generative AI and the expertise of physicians. Nat Biomed Eng. 2023; Available from: 10.1038/s41551-023-01160-9

16. Rodriguez P, Caccia M, Lacoste A, Zamparo L, Laradji I, Charlin L, et al. Beyond trivial counterfactual explanations with diverse valuable explanations. IEEE/CVF International Conference on Computer Vision (ICCV). 2021.

17. Sanchez P, Voisey JP, Xia T, Watson HI, O’Neil AQ, Tsaftaris SA. Causal machine learning for healthcare and precision medicine. R Soc Open Sci. 2022;9:220638.

18. Sanchez P, Kascenas A, Liu X, O’Neil AQ, Tsaftaris SA. What is Healthy? Generative Counterfactual Diffusion for Lesion Localization. Deep Generative Models. Springer Nature Switzerland; 2022. page 34–44.

19. Dolezal JM, Wolk R, Hieromnimon HM, Howard FM, Srisuwananukorn A, Karpeyev D, et al. Deep learning generates synthetic cancer histology for explainability and education. NPJ Precis Oncol. 2023;7:49.

20. Naglah A, Khalifa F, El-Baz A, Gondim D. Conditional GANs based system for fibrosis detection and quantification in Hematoxylin and Eosin whole slide images. Med Image Anal. 2022;81:102537.

21. Wölflein G, Um IH, Harrison DJ, Arandjelović O. HoechstGAN: Virtual lymphocyte staining using generative adversarial networks. Proceedings of the IEEE/CVF Winter Conference on Applications of Computer Vision. 2022. page 4997–5007.

22. Singla S, Eslami M, Pollack B, Wallace S, Batmanghelich K. Explaining the blackbox smoothly-A counterfactual approach. Med Image Anal. Elsevier BV; 2023;84:102721.

23. Lang O, Yaya-Stupp D, Traynis I, Cole-Lewis H, Bennett CR, Lyles CR, et al. Using generative AI to investigate medical imagery models and datasets. EBioMedicine. 2024;102:105075.

24. Howard FM, Hieromnimon HM, Ramesh S, Dolezal J, Kochanny S, Zhang Q, et al. Generative adversarial networks accurately reconstruct pan-cancer histology from pathologic, genomic, and radiographic latent features. Sci Adv. 2024;10:eadq0856.

25. Dhariwal P, Nichol A. Diffusion models beat GANs on image synthesis. Adv Neural Inf Process Syst. 2021;34:8780–94.

26. Song J, Meng C, Ermon S. Denoising Diffusion Implicit Models. International Conference on Learning Representations (ICLR). 2021.

27. Preechakul K, Chatthee N, Wizadwongsa S, Suwajanakorn S. Diffusion autoencoders: Toward a meaningful and decodable representation. Proc IEEE Comput Soc Conf Comput Vis Pattern Recognit. 2021;10609–19.

28. Meng C, He Y, Song Y, Song J, Wu J, Zhu J-Y, et al. SDEdit: Guided image synthesis and editing with stochastic differential equations. International Conference on Learning Representations (ICLR); 2022.

29. Augustine TN. Weakly-supervised deep learning models in computational pathology. EBioMedicine. Elsevier BV; 2022;81:104117.

30. Gadermayr M, Tschuchnig M. Multiple instance learning for digital pathology: A review of the state-of-the-art, limitations & future potential. Comput Med Imaging Graph. Elsevier BV; 2024;112:102337.

31. Kather JN, Pearson AT, Halama N, Jäger D, Krause J, Loosen SH, et al. Deep learning can predict microsatellite instability directly from histology in gastrointestinal cancer. Nat Med. Springer Science and Business Media LLC; 2019;25:1054– 6.

32. Kather, J. N., Halama, N., Marx, A. 100,000 histological images of human colorectal cancer and healthy tissue. 2018. Available from: https://zenodo.org/records/1214456

33. Macenko M, Niethammer M, Marron JS, Borland D, Woosley JT, Guan X, et al. A method for normalizing histology slides for quantitative analysis. 2009 IEEE International Symposium on Biomedical Imaging: From Nano to Macro. IEEE; 2009. page 1107–10.

34. Cortes-Ciriano I, Lee S, Park W-Y, Kim T-M, Park PJ. A molecular portrait of microsatellite instability across multiple cancers. Nat Commun. Springer Science and Business Media LLC; 2017;8:15180.

35. Komura D, Ishikawa S. Histology images from uniform tumor regions in TCGA Whole Slide Images. In Cell Reports (1.0, Vol. 38, Number 9, p. 110424). 2021. Available from: 10.5281/zenodo.5889558

36. Elfwing S, Uchibe E, Doya K. Sigmoid-weighted linear units for neural network function approximation in reinforcement learning. Neural Netw. Elsevier BV; 2018;107:3–11.

37. Wang Z, Simoncelli EP, Bovik AC. Multiscale structural similarity for image quality assessment. The Thirty-Seventh Asilomar Conference on Signals, Systems & Computers, 2003. IEEE; 2003. page 1398–402 Vol.2.

38. Wang Z, Bovik AC, Sheikh HR, Simoncelli EP. Image quality assessment: from error visibility to structural similarity. IEEE Trans Image Process. 2004;13:600– 12.

39. Parmar G, Zhang R, Zhu J-Y. On aliased resizing and surprising subtleties in GAN evaluation. Proc IEEE Comput Soc Conf Comput Vis Pattern Recognit. 2022;11400–10.

40. Vahadane A, Peng T, Sethi A, Albarqouni S, Wang L, Baust M, et al. Structure-preserving color normalization and sparse stain separation for histological images. IEEE Trans Med Imaging. Institute of Electrical and Electronics Engineers (IEEE); 2016;35:1962–71.

41. El Nahhas OSM, van Treeck M, Wölflein G, Unger M, Ligero M, Lenz T, et al. From whole-slide image to biomarker prediction: end-to-end weakly supervised deep learning in computational pathology. Nat Protoc. 2025;20:293–316.

42. Chen RJ, Ding T, Lu MY, Williamson DFK, Jaume G, Song AH, et al. Towards a general-purpose foundation model for computational pathology. Nat Med. 2024;30:850–62.

43. Zimmermann E, Vorontsov E, Viret J, Casson A, Zelechowski M, Shaikovski G, et al. Virchow2: Scaling self-supervised mixed magnification models in pathology. arXiv [cs.CV]. 2024. Available from: http://arxiv.org/abs/2408.00738

44. Lu MY, Chen B, Williamson DFK, Chen RJ, Liang I, Ding T, et al. A visual-language foundation model for computational pathology. Nat Med. 2024;30:863–74.

45. Ignatov A, Yates J, Boeva V. Histopathological image classification with cell morphology aware deep neural networks. Proceedings of the IEEE/CVF Conference on Computer Vision and Pattern Recognition. 2024. page 6913–25.

46. Cai Y, Lu Z, Chen C, Zhu Y, Chen Z, Wu Z, et al. An atlas of genetic effects on cellular composition of the tumor microenvironment. Nat Immunol. Springer Science and Business Media LLC; 2024;25:1959–75.

47. Hamilton SR. Carcinoma of the colon and rectum. Pathology and genetics of tumors of digestive system. 2000;

48. Travis WD. Pathology of lung cancer. Clin Chest Med. Elsevier BV; 2011;32:669– 92.

49. Schlageter M, Terracciano LM, D’Angelo S, Sorrentino P. Histopathology of hepatocellular carcinoma. World J Gastroenterol. Baishideng Publishing Group Inc.; 2014;20:15955–64.

50. Vijgen S, Terris B, Rubbia-Brandt L. Pathology of intrahepatic cholangiocarcinoma. Hepatobiliary Surg Nutr. AME Publishing Company; 2017;6:22–34.

51. Jenkins MA, Hayashi S, O’Shea A-M, Burgart LJ, Smyrk TC, Shimizu D, et al. Pathology features in Bethesda guidelines predict colorectal cancer microsatellite instability: a population-based study. Gastroenterology. Elsevier BV; 2007;133:48–56.

52. Shia J, Schultz N, Kuk D, Vakiani E, Middha S, Segal NH, et al. Morphological characterization of colorectal cancers in The Cancer Genome Atlas reveals distinct morphology-molecular associations: clinical and biological implications. Mod Pathol. 2017;30:599–609.

53. Puccini A, Poorman K, Catalano F, Seeber A, Goldberg RM, Salem ME, et al. Molecular profiling of signet-ring-cell carcinoma (SRCC) from the stomach and colon reveals potential new therapeutic targets. Oncogene. Springer Science and Business Media LLC; 2022;41:3455–60.

54. Reitsam NG, Märkl B, Dintner S, Waidhauser J, Vlasenko D, Grosser B. Concurrent loss of MLH1, PMS2 and MSH6 immunoexpression in digestive system cancers indicating a widespread dysregulation in DNA repair processes. Front Oncol. Frontiers Media SA; 2022;12:1019798.

55. Remo A, Fassan M, Vanoli A, Bonetti LR, Barresi V, Tatangelo F, et al. Morphology and molecular features of rare colorectal carcinoma histotypes. Cancers (Basel). MDPI AG; 2019;11:1036.

56. Howard FM, Dolezal J, Kochanny S, Schulte J, Chen H, Heij L, et al. The impact of site-specific digital histology signatures on deep learning model accuracy and bias. Nat Commun. Springer Science and Business Media LLC; 2021;12:4423.

57. Dehkharghanian T, Bidgoli AA, Riasatian A, Mazaheri P, Campbell CJV, Pantanowitz L, et al. Biased data, biased AI: deep networks predict the acquisition site of TCGA images. Diagn Pathol. 2023;18:67.

58. Kheiri F, Rahnamayan S, Makrehchi M, Asilian Bidgoli A. Investigation on potential bias factors in histopathology datasets. Sci Rep. Nature Publishing Group; 2025;15:11349.

