## Supplementary Material for "Counterfactual Diffusion Models for Interpretable Explanations of Artificial Intelligence Models in Pathology"

Laura Žigutytė, Tim Lenz, Tianyu Han, Katherine Jane Hewitt, Nic Gabriel Reitsam, Sebastian Foersch, Zunamys Itzell Carrero, Michaela Unger, Asier Rabasco Meneghetti, Alexander T. Pearson, Daniel Truhn, Jakob Nikolas Kather

### Supplementary Figures

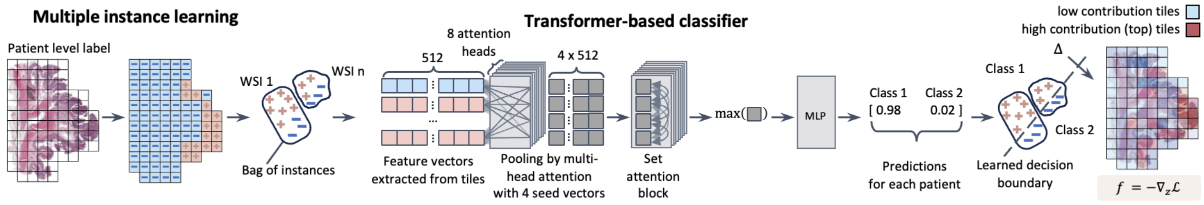


**Figure S1. End-to-end multiple-instance learning (MIL) with a transformer for patient-level histopathology classification and tile-level interpretability.** A whole-slide image (WSI) is partitioned into fixed-size tiles (e.g., 512 x 512 px at 0.5 micrometers per pixel for TCGA-CRC). Tiles from one or more WSIs of the same patient are grouped into a bag of instances and inherit the patient-level label (weak supervision; no per-tile labels). Each tile is encoded into a 512-dimensional feature vector using the pretrained MoPaDi encoder. Features are processed by the transformer: pooled multi-head attention (PMA) with 4 seed vectors and 8 heads aggregates the unordered set into a small set of pooled descriptors; a set attention block refines interactions among pooled descriptors. The architecture is permutation-invariant by design. A layer-norm precedes pooling. The pooled descriptors are collapsed to a single patient embedding via element-wise max over seeds. The patient embedding passes through a two-layer multilayer perceptron (MLP) to produce logits and class probabilities. The learned decision boundary separates patient bags in embedding space; the predicted class and confidence are shown. To identify tiles most influential for the model's decision, each tile feature vector is independently evaluated as a single-instance "bag", yielding a per-tile class probability. Tiles are ranked by their individual softmax probability for the predicted class, and the highest-scoring ("top") tiles are highlighted as high-contribution regions, while lower-scoring tiles are shown as low-contribution regions. This produces a tile-level importance map reflecting the tiles whose isolated features most strongly support the patient-level prediction, complementing the weakly-supervised MIL framework. Top-contributing tiles are subsequently used for counterfactual image generation.

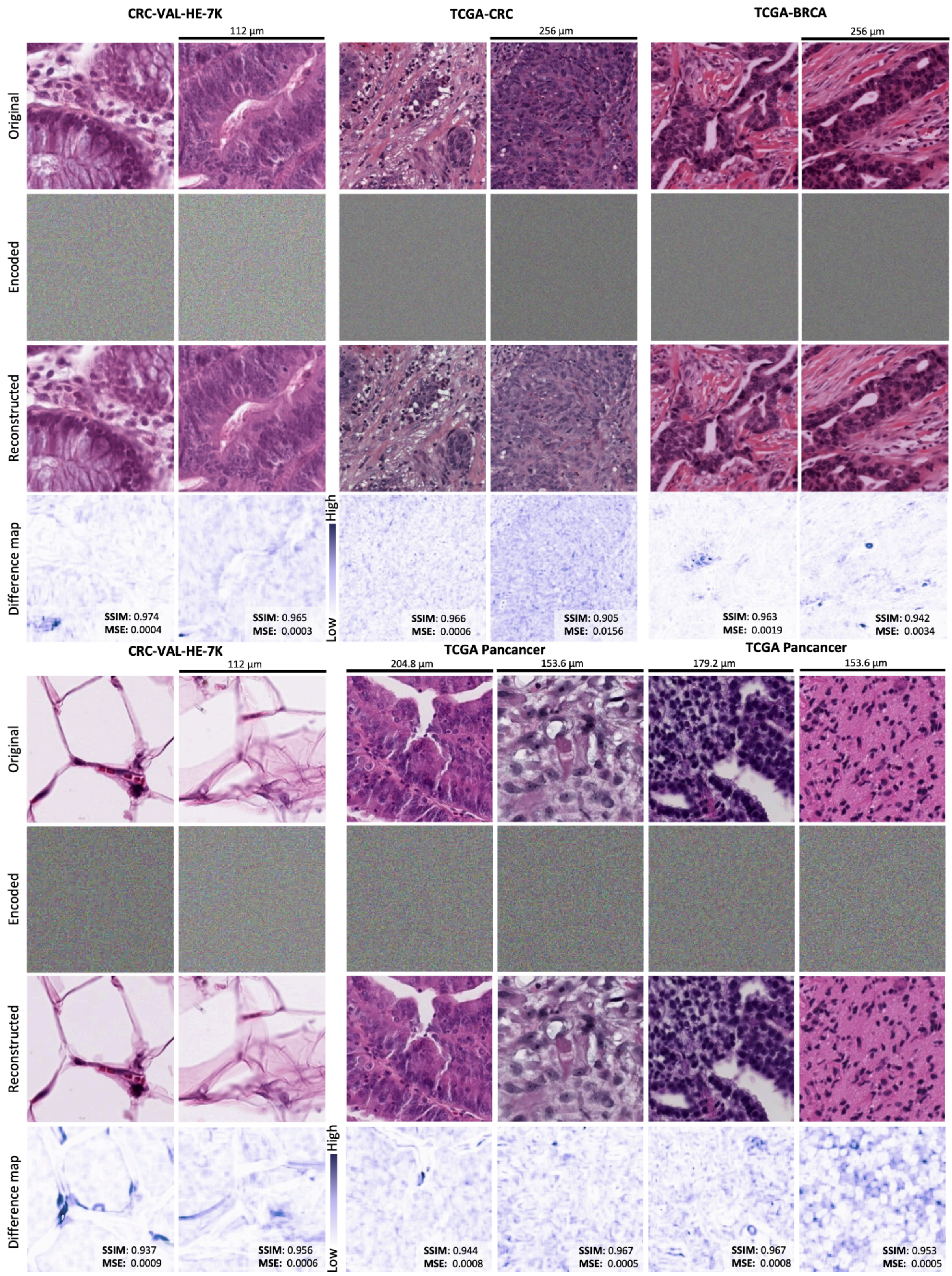


**Figure S2. Representative examples for all datasets of encoded images into noise and reconstructed back to the original.** Random representative tiles from each dataset are followed by encoded images into noise with a custom-trained diffusion autoencoder, and reconstructed images back to the original ones. The model sometimes struggles with structure boundaries and white spaces. Difference images visually represent dissimilarity between the original images and reconstructed images (white regions indicate perfect similarity, while darker regions indicate dissimilarity) and are displayed together with quantitative metrics for the reconstruction quality. The scale bar, if not indicated otherwise, corresponds to all images within one panel. MSE, mean square error; SSIM, structural similarity index measure.

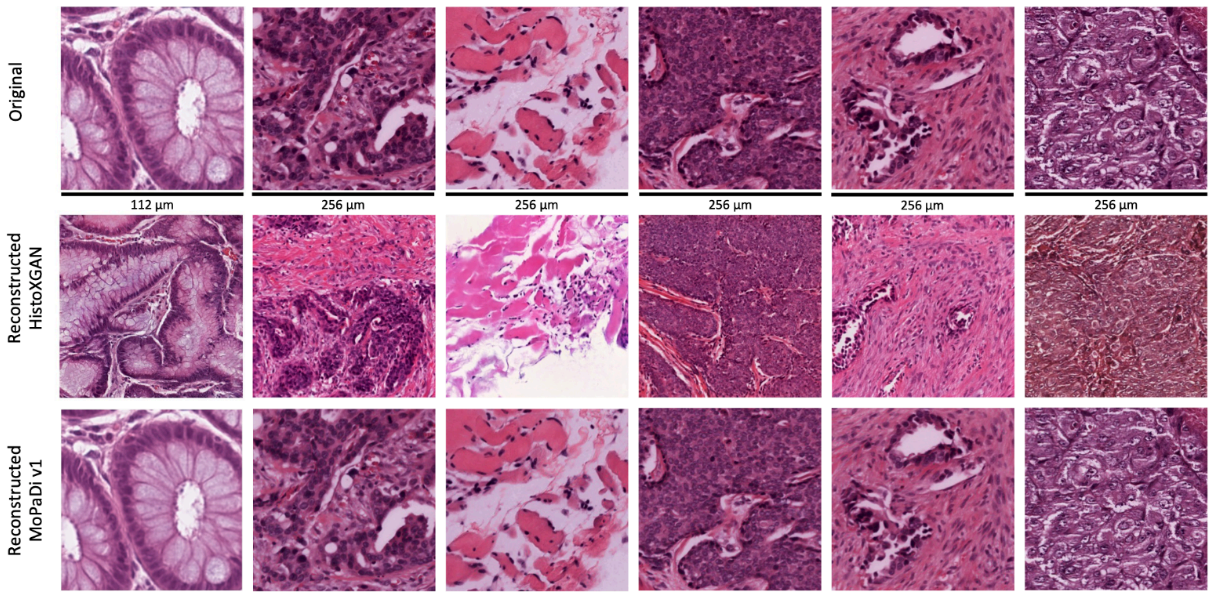


**Figure S3. Qualitative comparison of MoPaDi's reconstructions with one of the GAN-based approaches (HistoXGAN).** Due to the addition of a stochastic noise map in MoPaDi, the structural details of the original images are better preserved in the reconstructions.

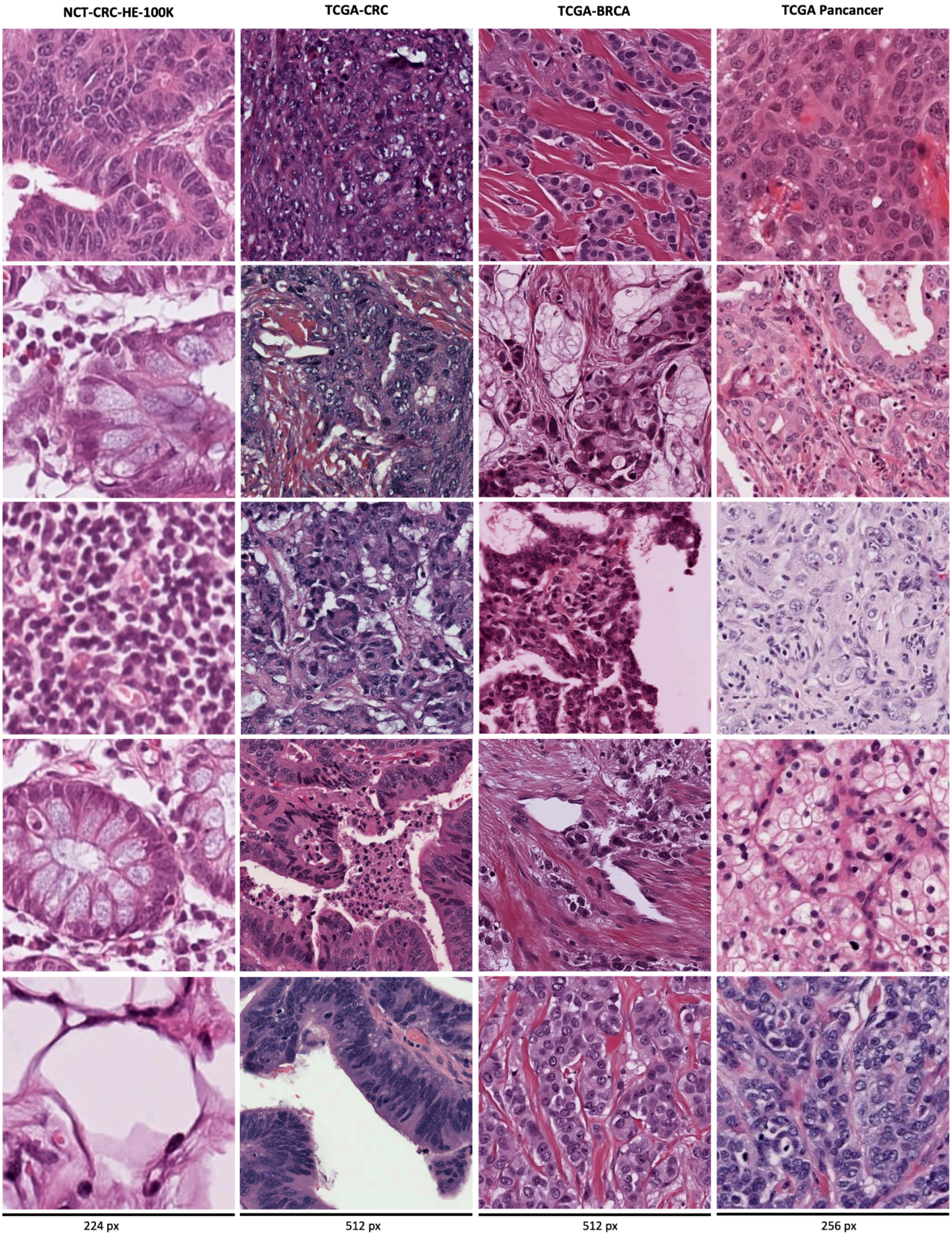


**Figure S4. Randomly sampled representative examples of generated synthetic images with four trained models** trained on different datasets: NCT-CRC-HE-100K, TCGA-CRC, TCGA-BRCA, and TCGA Pan-cancer. The scale bar corresponds to the image size the generative model was trained on.

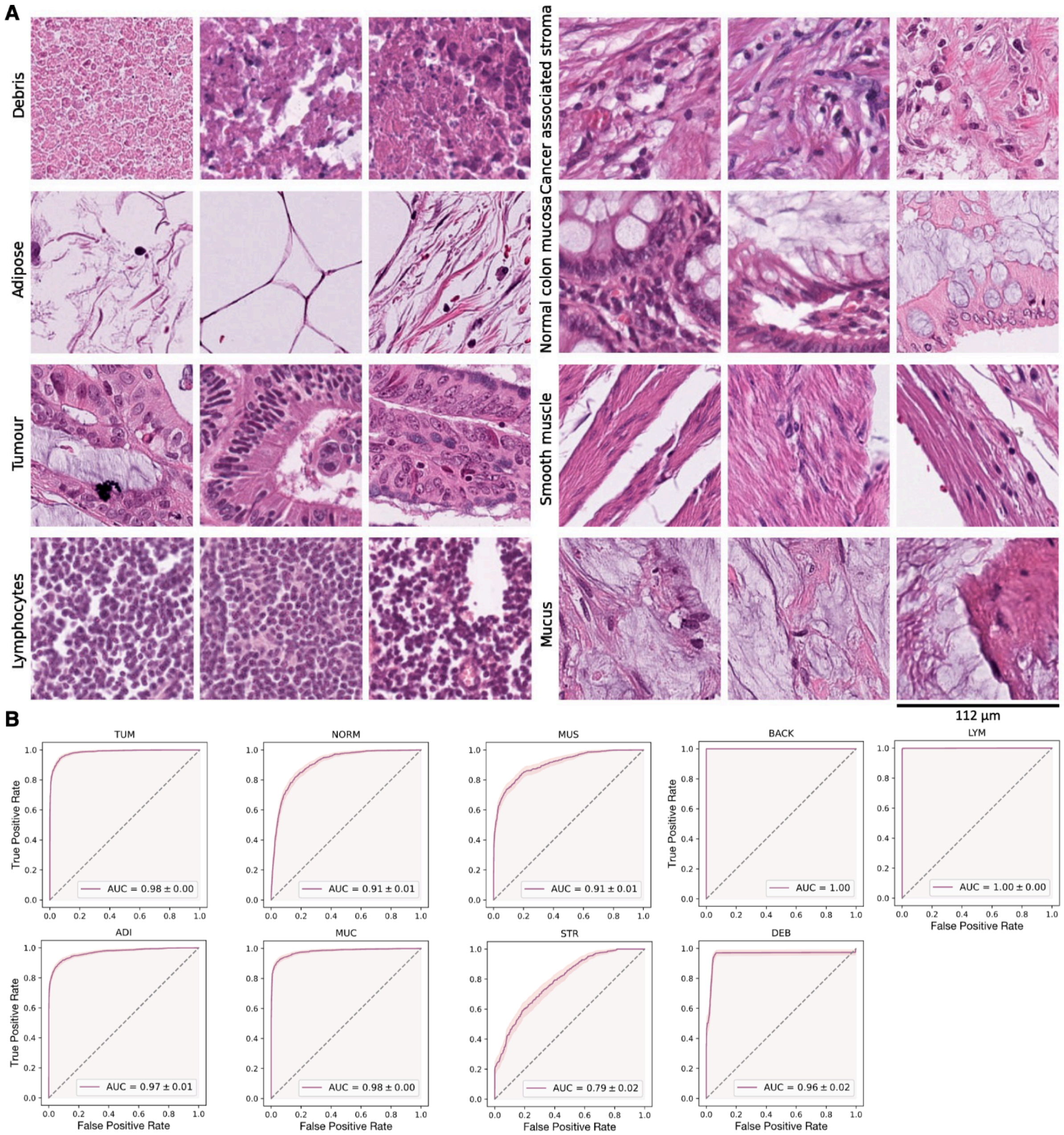


**Figure S5. Representative examples of classes in the NCT-CRC-HE-100K dataset and receiver operating characteristic curves (ROC) evaluating the trained classifier's performance on the CRC-VAL-HE-7K dataset (*N* = 7,180 tiles). A,** NCT-CRC-HE-100K dataset contains nine classes: debris (DEB), adipose (ADI), tumor (i.e., dysplastic epithelium; TUM), lymphocytes (LYM), cancer-associated stroma (STR), normal colon mucosa (NORM), smooth muscle (MUS), mucus (MUC), and background (not displayed; BACK). **B,** Evaluation of the trained classifier's performance on the CRC-VAL-HE-7K shows good performance, with an area under the ROCs ranging from 0.79 ± 0.02 for the STR class to 1.00 ± 0.00 for the BACK and LYM classes. AUC, area under the curve.

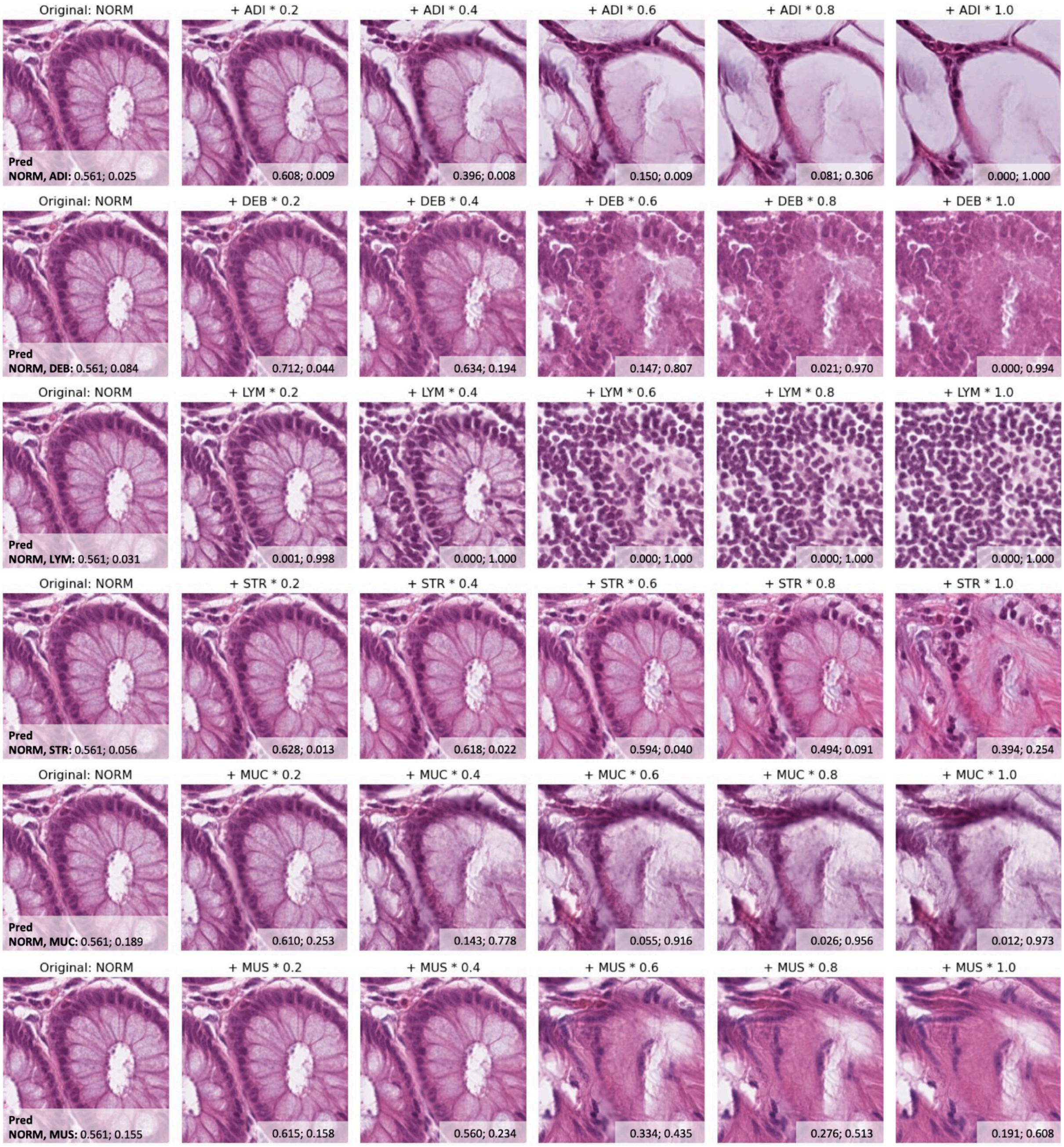


**Figure S6. Examples of counterfactual explanations for the tissue type classifier**, showing how the original histology tile would need to appear to be reassigned to the indicated target class. Each row contains (**left**) the original normal colon mucosa (NORM) tile and shows (**right**) images obtained after progressively larger latent-space shifts toward a target class (manipulation amplitude *α* sampled at increments of 0.2 from 0.2 to 1.0). The numbers on every tile are the sigmoid scores for two classes. For five classes (adipose [ADI], debris [DEB], lymphocytes [LYM], mucus [MUC], smooth muscle [MUS]), the model's prediction flips to the target when $\alpha \geq 0.6$. In contrast, the cancer-associated stroma (STR) counterfactual images never cross the decision boundary. This suggests that STR tiles lie farther from NORM in latent space, which likely contributes to the model's poorer classification performance on STR (see **Fig. S5B** for performance metrics).

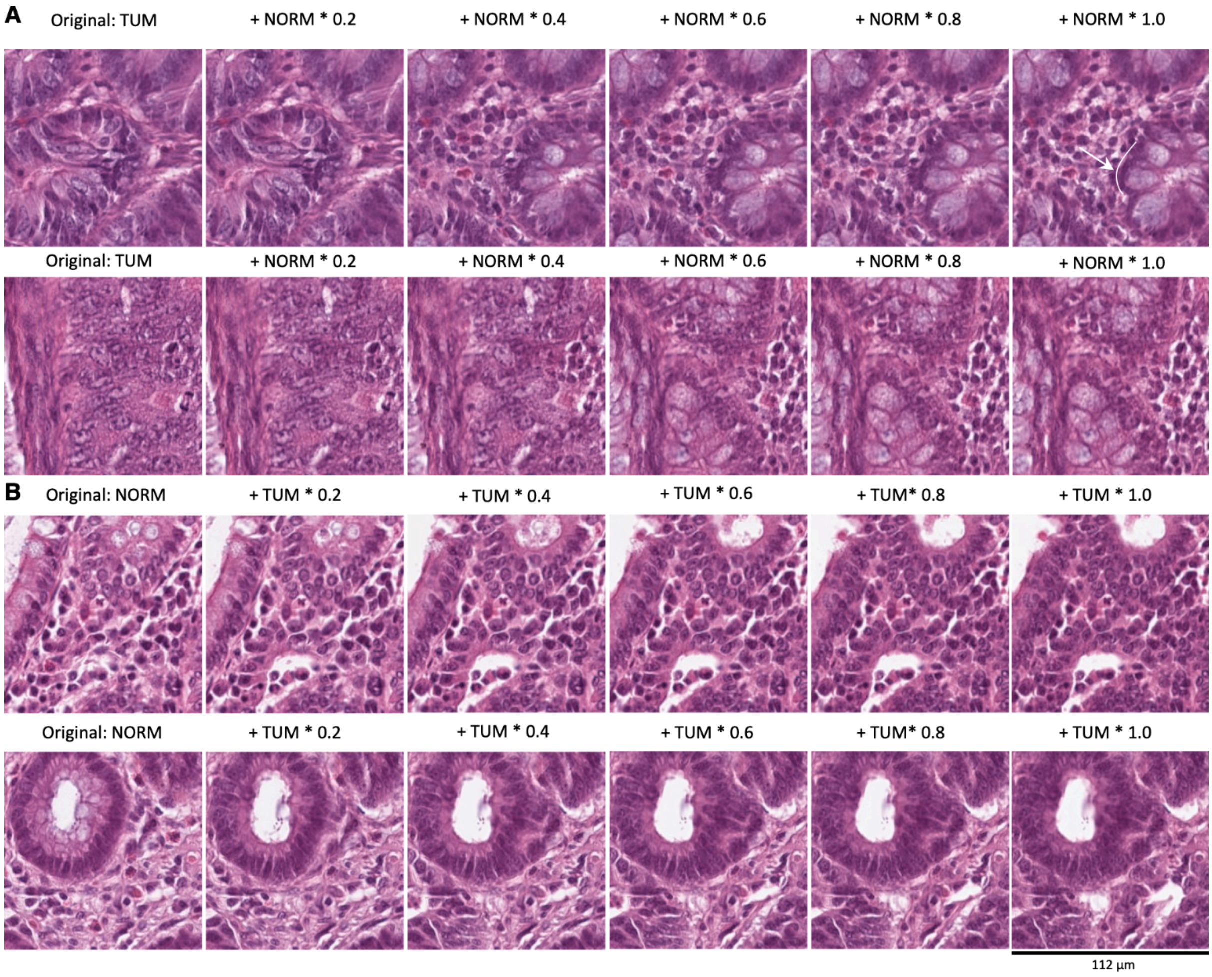


**Figure S7. Representative examples of transitions to counterfactual images for the CRC-VAL-HE-7K dataset.** These examples were included in a user study with two pathologists. In some cases, subtle features, such as insufficient hyperchromasia or gland formation without a visible basement membrane, resulting in unrealistically smooth boundaries between cells and adjacent stroma, hinted at the images' synthetic origin. The manipulation amplitude (*α*) is given above each image. (**A**) Dysplastic epithelium tile (TUM) manipulated to healthy colon mucosa (NORM). The model captured the appearance of glands; however, no basement membrane (white arrow) was generated in both examples. (**B**) Healthy colon mucosa tile manipulated to dysplastic epithelium. Nuclei in the top synthetic counterfactual image lacked hyperchromatism. Such patterns may reflect the model's internal prioritization of morphological features; for example, the presence of goblet cells might contribute more strongly to class prediction than the delineation of gland boundaries.

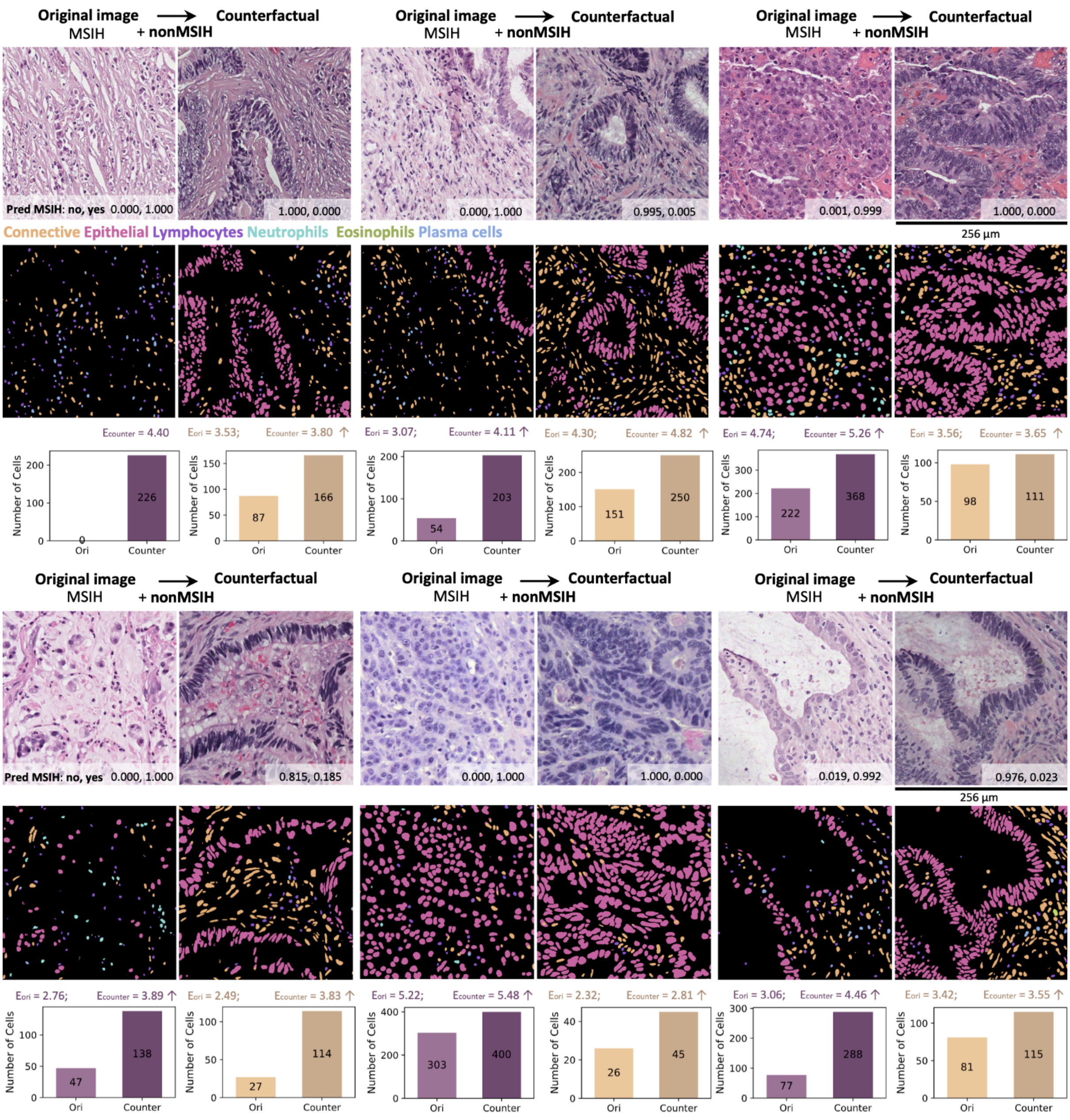


**Figure S8. Additional examples of generated counterfactual images for tiles from patients with high microsatellite instability (MSIH).** Representative tiles from MSIH patients with corresponding cell segmentation maps showing distinct cell types (connective tissue cells in orange, epithelial cells in pink, lymphocytes in purple, eosinophils in green, and plasma cells in blue) are depicted in two rows. For each case, original MSIH images and their counterfactuals, i.e., images manipulated to be classified as nonMSIH, are compared. Entropy values ($E$) are shown for each image, with arrows indicating an increase (↑) or a decrease (↓) in counterfactual images. The counterfactual images exhibit consistent patterns: an increase in epithelial and connective tissue cell count coupled with higher spatial entropy. Furthermore, a decrease in plasma cells and lymphocytes can be observed qualitatively. Epithelial cells are arranged in a more glandular pattern. The scale bar, 256 µm, applies to all tiles.

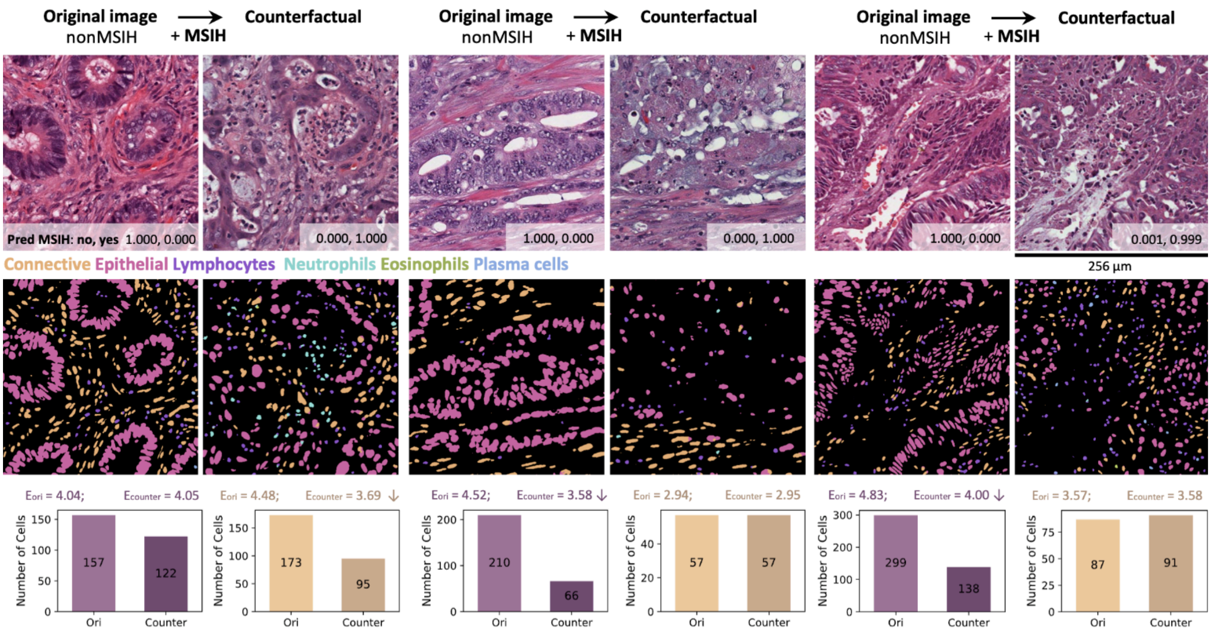


**Figure S9. Additional examples of generated counterfactual images for tiles from microsatellite stable (nonMSIH) patients.** The top row contains representative tiles from nonMSIH patients; (**middle**) corresponding cell segmentation maps showing distinct cell types (connective tissue cells in orange, epithelial cells in pink, lymphocytes in purple, eosinophils in green, and plasma cells in blue). For each case, original nonMSIH images and their counterfactuals, i.e., images manipulated to be classified as MSIH, are compared. Entropy values (*E*) are shown for each image, with arrows indicating an increase (↑) or a decrease (↓) in counterfactual images. In contrast to the examples in **Fig. S8**, the counterfactual images show a decreased epithelial cell count along with lower spatial entropy. The scale bar, 256 µm, applies to all tiles.

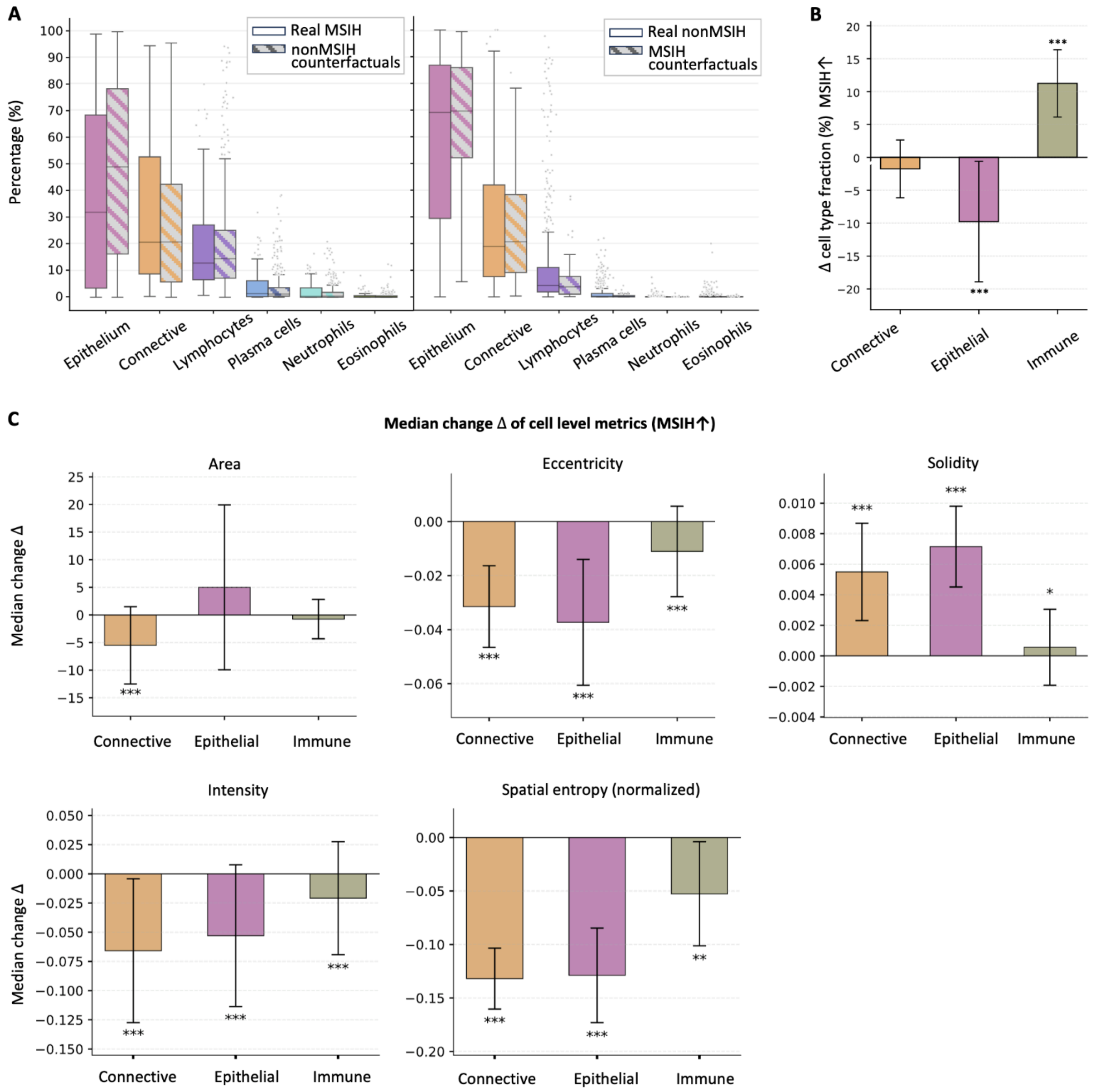


**Figure S10. Cellular composition and morphological alterations associated with MSIH counterfactual transitions in TCGA CRC.** (**A**) The distribution of cell-type percentages in: (**left**) real MSIH ($N$ = 65) and nonMSIH counterfactual tiles ($N$ = 475), (**right**): real nonMSIH tiles ($N$ = 475) and counterfactual tiles ($N$ = 65). Boxes/whiskers summarize tile-level values; gray dots are individual outlier tiles. Statistical inference was performed at the patient level (per-patient mean percentage). No cell type showed a statistically significant difference after multiple-testing correction across six tests; full statistics ($p$, adjusted $p$, Cliff's δ with 95% CI) are reported in **Table S4**. (**B**) Change in cell-type fractions across connective, epithelial, and immune cell types when manipulating toward MSIH. MSIH counterfactuals show a significant increase in immune and connective tissue and a reduction in epithelial fraction ($p$ < 0.001, Wilcoxon paired test). (**C**) Median changes (Δ) in cell-level morphological metrics for counterfactual manipulations toward MSIH.

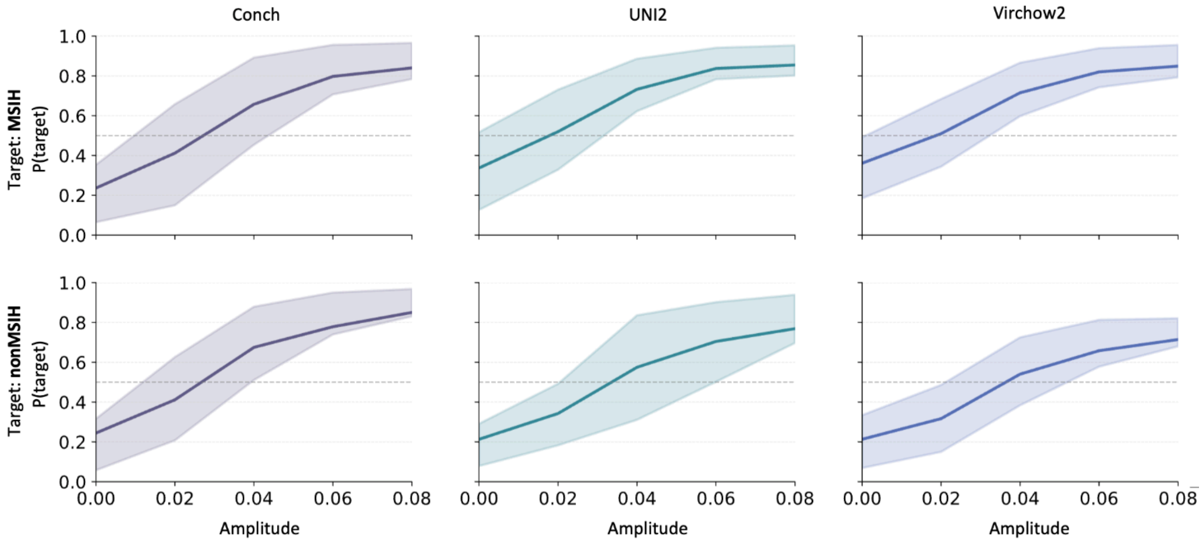


**Figure S11. Validation of counterfactual transitions using independent histopathology classifiers.** Prediction probabilities across counterfactual manipulation amplitudes ($\alpha$) for three independent foundation model-based MSI classifiers (Conch, UNI2, and Virchow2). Lines represent the mean predicted probability for the target class, and shaded regions denote the interquartile range across tiles. Across all models and directions, confidence shifts monotonically toward the intended class, demonstrating that MoPaDi counterfactual trajectories are supported by external classifiers and reflect biologically meaningful decision directions rather than model-specific artifacts.

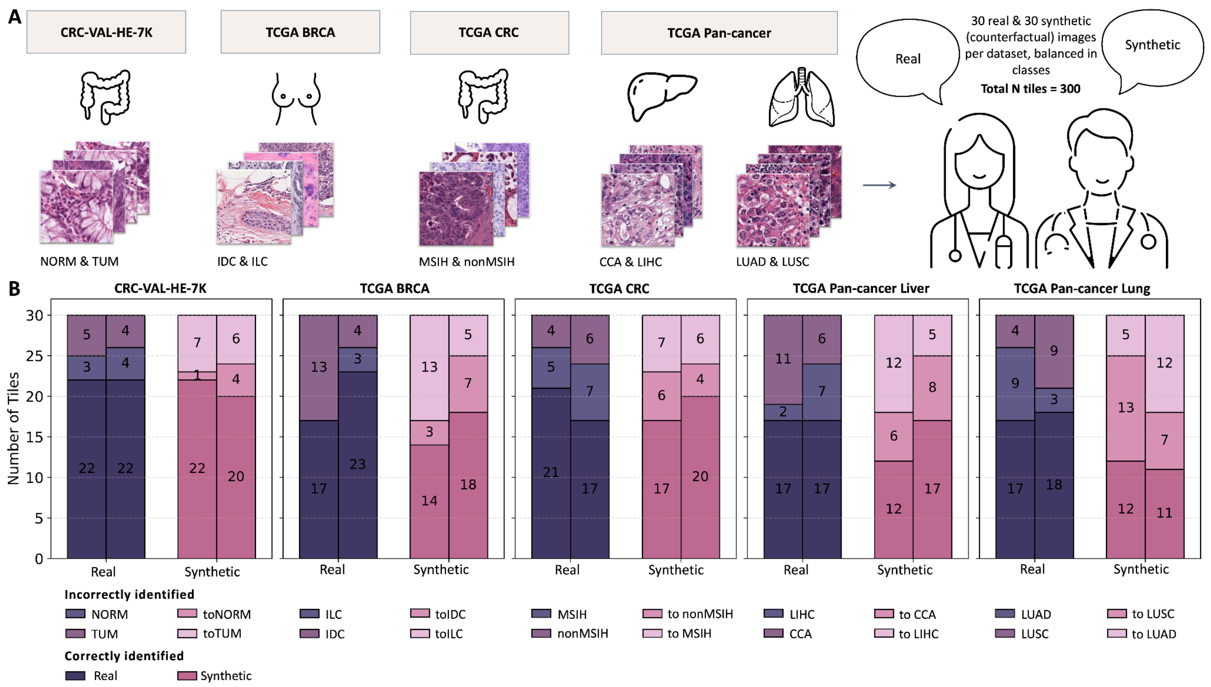


**Figure S12. Overview and results of the tile-level user study evaluating real vs. synthetic histopathology images to further assess the perceptual quality of counterfactual images.** (**A**) For each dataset (CRC-VAL-HE-7K, TCGA-BRCA, TCGA-CRC, and two tasks for the TCGA Pan-cancer dataset), 30 real tiles (15 per class) and 30 corresponding synthetic (counterfactual) tiles were randomly selected (Total $N$ = 300). Two pathologists independently reviewed the tiles and classified each one as either real or synthetic. (**B**) Classification performance of both pathologists (indicated by separate columns) across tasks, separated into real and synthetic tiles. Bars show the number of tiles per category: correctly identified (dark), or incorrectly classified with a ground truth class indication (lighter shades). Across all tasks, pathologists were more accurate in identifying real tiles than synthetic ones. Pathologists misclassified between 26.7% and 63.3% of synthetic images as real across all tasks, indicating that MoPaDi-generated images often exhibit high perceptual realism. NORM, healthy colon mucosa; TUM, colorectal adenocarcinoma epithelium; IDC, invasive ductal carcinoma; ILC, invasive lobular carcinoma; CCA, cholangiocarcinoma; MSIH, microsatellite instability-high; nonMSIH, microsatellite stable; LIHC, liver hepatocellular carcinoma; LUSC, lung squamous cell carcinoma; LUAD, lung adenocarcinoma.

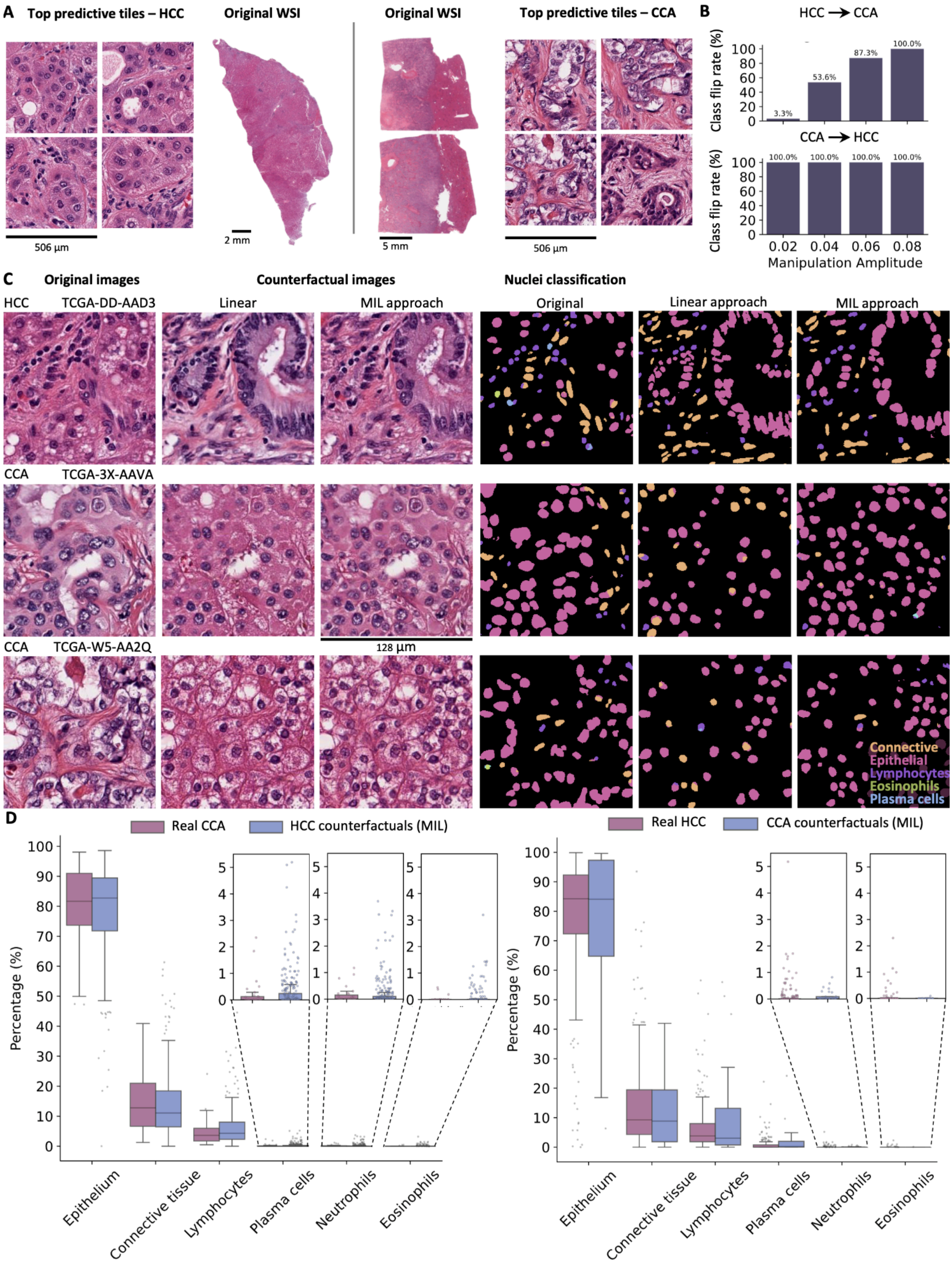


**Figure S13. Comparison of counterfactual images obtained with a linear approach and multiple instance learning (MIL) for the liver cancer type classification.** (**A**) Top predictive tiles for hepatocellular carcinoma (HCC) and cholangiocarcinoma (CCA) of the MIL-based classifier. (**B**) Counterfactual image generation effectiveness, measured as the percentage of generated images, across varying manipulation amplitudes, predicted as the opposite class. HCC to CCA required higher amplitudes to reach similar results, which likely reflects class imbalance in the dataset. (**C**) Representative examples of generated counterfactual images using both approaches and a comparison of the obtained segmentation masks. (**D**) The distribution of different cell types in real and synthetic tiles. (**Left**) CCA tiles (31 tiles from 6 patients) and synthetic (counterfactual images for HCC class, i.e., the most manipulated tiles, 276 tiles from 54 patients). No significant differences in cell type percentages between the groups were observed after applying the Bonferroni correction for multiple comparisons, based on a two-sided Mann-Whitney U test (adjusted $p > 0.05$). (**Right**) Real HCC tiles (276 tiles from 54 patients) vs. synthetic (counterfactual images for CCA class, i.e., the most manipulated tiles, 31 tiles from 6 patients). No significant differences in cell type percentages between groups were observed. A full description of the statistical parameters, including *p*-values, is provided in **Table S5**. AP, average precision.

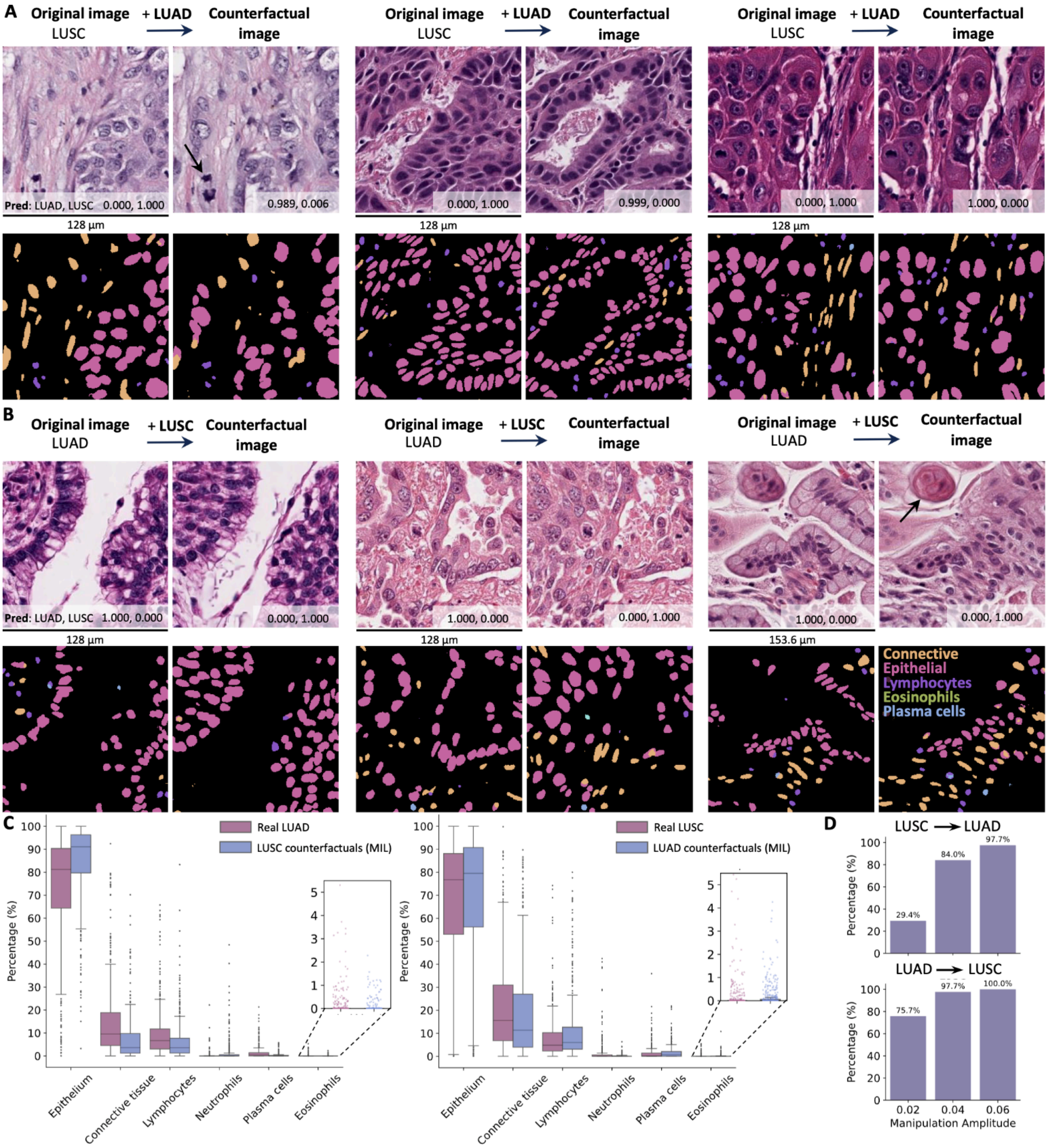


**Figure S14. Results of counterfactual image generation for multiple instance learning (MIL)-based lung cancer type classifier.** (**A**) Representative examples of lung squamous cell carcinoma (LUSC) to lung adenocarcinoma (LUAD) transitions to counterfactual images. (**Left**) black arrow – remaining mitotic figure, which is rather a typical feature of LUSC than LUAD; (**middle**) glands with nuclear molding, which was a prominent feature in the original image and the counterfactual image as well; (**right**) remaining visible squamous differentiation in the counterfactual image with the solid morphology. (**B**) Representative examples of LUAD to LUSC transitions to counterfactual images. (**Left)** Remaining glandular architecture with more cell layers in the counterfactual image; (**middle**) potentially visible intercellular bridges in the counterfactual image and a higher degree of nuclear atypia; (**right**) black arrow –

potentially visible squamous pearl. (**C**) The distribution of different cell types in real and synthetic tiles. (**Left**) Real LUAD tiles ($N$ = 470) and synthetic (counterfactual images of LUSC, i.e., the most manipulated tiles [$\alpha$ = 0.06], where 98.8% of counterfactual images were predicted as the target class, $N$ = 470). Statistically ($p <$ 0.01) and practically (Cliff's delta < 0.147) significant differences in cell type percentages were observed for all groups, except eosinophils. (**Right**) Real LUSC tiles ($N$ = 470) vs. LUAD counterfactual images ($N$ = 470). Statistically significant differences in cell type percentages were observed for neutrophils and plasma cells. However, these differences were practically significant only for plasma cells. A full description of the statistical parameters, including $p$-values, is provided in **Table S6**. (**D**) Quantitative evaluation of counterfactual image generation effectiveness. Bars show the percentage of successful manipulations (i.e., predicted class equals target class) across different manipulation amplitudes.

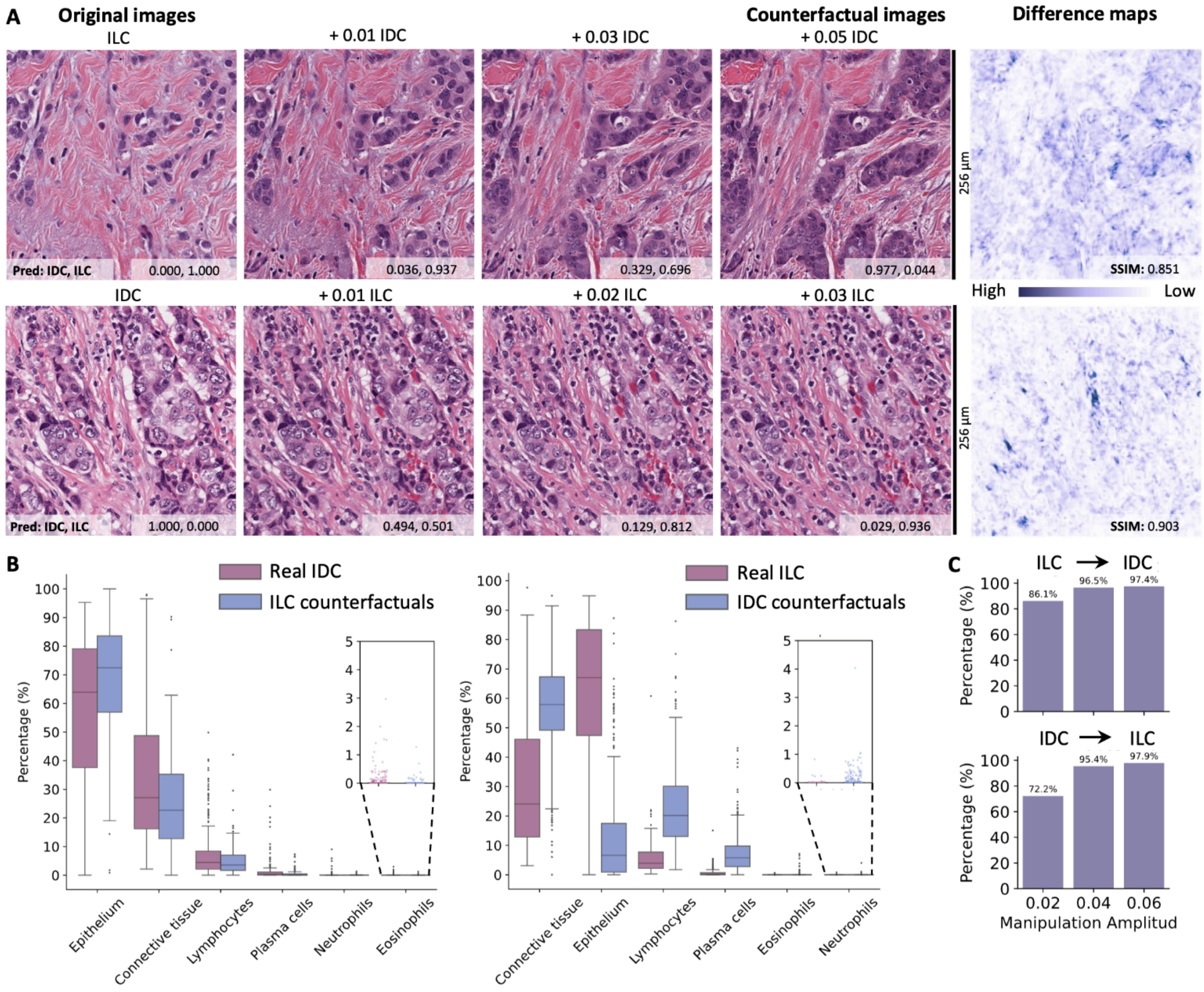


**Figure S15. Evaluation of breast cancer types multiple instance learning (MIL)-based classifier (invasive ductal carcinoma [IDC] vs. invasive lobular carcinoma [ILC]), and generated counterfactual images.** (**A**) Representative examples of invasive lobular carcinoma (ILC) tile transitioning to its counterfactual invasive ductal carcinoma (IDC) image and vice versa. Difference maps display pixel-wise differences between the original and the synthetic tile. SSIM, structural similarity index measure. (**B**) The distribution of different cell types in real and synthetic images. (**Left**) IDC tiles ($N$ = 475) vs. synthetic (counterfactual images, i.e., the most manipulated tiles [$\alpha$ = 0.06], where 97.4% of counterfactual images were predicted as the target class; $N$ = 115). Statistically ($p$ < 0.01) and practically (Cliff's delta < 0.147) significant differences in cell type percentages were observed for epithelial and plasma cells. (**Right**) Real IDC ($N$ = 115) and synthetic ($N$ = 475) tiles. Statistically significant differences in cell type percentages were observed for all cell types, except neutrophils and eosinophils. However, these differences were practically significant only for plasma cells. A full description of the statistical parameters, including $p$-values, is provided in **Table S7**. (**C**) Quantitative evaluation of counterfactual image generation effectiveness. Bars show the percentage of successful manipulations (i.e., predicted class equals target class) across different manipulation amplitudes.

### Supplementary Tables

**Table S1. Patient characteristics of the TCGA-CRC cohort.** Patients without available whole slide images, missing resolution value in the metadata (*N* = 26), or available tumor annotations (*N* = 8) were excluded. WSIs, whole slide images.

| Characteristic | | Total *N* patients = 582 | Train *N* = 474 (479 WSIs) | Test *N* = 108 (109 WSIs) |
|---|---|---|---|---|
| Age (years), median [IQR] | | 68.0 [58.0, 75.2] | 68.0 [58.0, 76.0] | 65.0 [57.0, 75.0] |
| | Missing | 2 | 2 | 1 |
| Follow-up (days), median [IQR] | | 555.0 [328.0, 1054.0] | 617.0 [329.5, 1088.5] | 521.0 [321.5, 925.5] |
| | Missing | 161 | 160 | 1 |
| Sex, n (%) | Male | 302 (51.9%) | 245 (51.7%) | 57 (52.8%) |
| | Female | 278 (47.8%) | 227 (47.9%) | 51 (47.2%) |
| | Missing | 2 (0.3%) | 2 (0.4%) | 0 |
| Organ, n (%) | Colon | 319 (54.8%) | 233 (49.2%) | 86 (79.6%) |
| | Rectum | 104 (17.9%) | 83 (17.5%) | 21 (19.4%) |
| | Missing | 159 (27.3%) | 158 (33.3%) | 1 (0.9%) |
| Stage, n (%) | 1.0 | 69 (11.9%) | 52 (11.0%) | 17 (15.7%) |
| | 2.0 | 150 (25.8%) | 106 (22.4%) | 44 (40.7%) |
| | 3.0 | 134 (23.0%) | 106 (22.4%) | 28 (25.9%) |
| | 4.0 | 57 (9.8%) | 42 (8.9%) | 15 (13.9%) |
| | Missing | 172 (29.6%) | 168 (35.4%) | 4 (3.7%) |
| Status, n (%) | Alive | 335 (57.6%) | 246 (51.9%) | 89 (82.4%) |
| | Dead | 86 (14.8%) | 68 (14.3%) | 18 (16.7%) |
| | Missing | 161 (27.7%) | 160 (33.8%) | 1 (0.9%) |
| Laterality, n (%) | Left | 225 (38.7%) | 173 (36.5%) | 52 (48.1%) |
| | Right | 167 (28.7%) | 124 (26.2%) | 43 (39.8%) |
| | Missing | 190 (32.6%) | 177 (37.3%) | 13 (12.0%) |
| BRAF, n (%) | WT | 437 (75.1%) | 339 (71.5%) | 98 (90.7%) |
| | MUT | 58 (10.0%) | 48 (10.1%) | 10 (9.3%) |
| | Missing | 87 (14.9%) | 87 (18.4%) | 0 |
| KRAS, n (%) | WT | 289 (49.7%) | 235 (49.6%) | 54 (50.0%) |
| | MUT | 206 (35.4%) | 152 (32.1%) | 54 (50.0%) |
| | Missing | 87 (14.9%) | 87 (18.4%) | 0 |
| MSI status, n (%) | nonMSIH | 360 (61.9%) | 266 (56.1%) | 94 (87.0%) |
| | MSIH | 61 (10.5%) | 48 (10.1%) | 13 (12.0%) |
| | Missing | 161 (27.7%) | 160 (33.8%) | 1 (0.9%) |

**Table S2. Distribution of patients in training and testing splits.** Numbers in brackets indicate the number of WSIs, accounting for cases where patients have more than one slide available. During the training of a diffusion autoencoder, the 'total train' number of patients was used, while during the training of the classifier, patients with unknown/other classes were ignored. LUAD, lung adenocarcinoma; LUSC, lung squamous cell carcinoma; HCC, hepatocellular carcinoma; CCA, cholangiocarcinoma; BRCA, breast invasive carcinoma; ILC, invasive lobular carcinoma; IDC, invasive ductal carcinoma; NaN, unknown.

| | **Lung cancer types** | | **Liver cancer types** | | **TCGA BRCA** | | | |
|---|---|---|---|---|---|---|---|---|
| | **LUAD** | **LUSC** | **HCC** | **CCA** | **ILC** | **IDC** | **Other** | **NaN** |
| **Train** | 348 | 350 | 218 | 26 | 108 | 416 | 169 | 128 |
| **Test** | 86 | 87 | 54 | 6 | 23 | 95 | 36 | 71 |
| **Total train** | 698 (775) | | 244 (247) | | 821(881) | | | |
| **Total test** | 173 (188) | | 60 (61) | | 225 (234) | | | |

**Table S3. Statistical characteristics of cell type percentages across groups: original and counterfactual images of two tissue types (malignant epithelium and normal colon mucosa).** For each cell type, the following metrics are reported: mean (%) ± standard deviation (SD); median (%) and the interquartile range (IQR, %); Hodges-Lehmann (HL) median shift in percentage points with 95% confidence intervals (CI); raw *p* value (*p* raw); corrected *p* value (*p* corr) after Bonferroni correction for multiple comparisons. After correction: *** $p < 0.001$; ** $p < 0.01$; * $p < 0.05$; *ns* – not significant. Qualitative interpretation of Cliff's delta (δ) to supplement HL shift: |δ| < 0.147 = negligible (*ng*); < 0.33 = small *; < 0.474 = medium **; ≥ 0.474 = large ***. TUM, malignant epithelium; NORM, normal colon mucosa; Manip to TUM, counterfactual images of NORM (synthetic); Manip to NORM, counterfactual images of TUM (synthetic).

| Cell type | Image type | Mean ± SD | Median | IQR | HL shift (95% CI), pp | *p* raw | *p* corr |
|---|---|---|---|---|---|---|---|
| **Epithelial** | TUM Original | 94.667 ± 7.192 | 97.186 | 5.744 | -1.57 (-1.83, -1.34) ** | $1.57e^{-58}$ | $9.44e^{-58}$ *** |
| | Manip to TUM | 98.367 ± 2.184 | 99.024 | 1.628 | | | |
| | NORM Original | 61.911 ± 17.395 | 63.462 | 21.158 | -2.47 (-4.04, -0.82) *ng* | $3.19e^{-3}$ | $1.91e^{-3}$ * |
| | Manip to NORM | 63.940 ± 18.906 | 65.868 | 25.341 | | | |
| **Connective** | TUM Original | 3.705 ± 5.415 | 1.808 | 4.177 | 1.27 (1.07, 1.47) *** | $5.67e^{-78}$ | $3.40e^{-77}$ *** |
| | Manip to TUM | 0.674 ± 1.063 | 0.306 | 0.922 | | | |
| | NORM Original | 15.606 ± 8.773 | 13.731 | 10.330 | 3.04 (2.38, 3.73) * | $1.81e^{-19}$ | $1.08e^{-18}$ *** |
| | Manip to NORM | 12.211 ± 7.322 | 11.249 | 9.086 | | | |
| **Lymphocytes** | TUM Original | 1.330 ± 2.328 | 0.637 | 1.356 | 0.18 (0.11, 0.24) * | $1.60e^{-10}$ | $9.57e^{-10}$ *** |
| | Manip to TUM | 0.773 ± 1.265 | 0.361 | 0.899 | | | |
| | NORM Original | 12.963 ± 8.312 | 10.646 | 11.051 | -2.27 (-2.94, -1.51) * | $9.98e^{-10}$ | $5.99e^{-9}$ *** |
| | Manip to NORM | 15.575 ± 9.840 | 13.760 | 10.870 | | | |
| **Plasma cells** | TUM Original | 0.153 ± 0.712 | 0.000 | 0.000 | 0.00 (0.00, 0.00) *ng* | $4.04e^{-4}$ | $2.43e^{-3}$ ** |
| | Manip to TUM | 0.129 ± 0.399 | 0.000 | 0.016 | | | |
| | NORM Original | 7.853 ± 6.315 | 6.380 | 7.498 | 0.47 (0.00, 0.97) *ng* | 0.042 | 0.251 *ns* |
| | Manip to NORM | 7.037 ± 5.380 | 5.989 | 7.454 | | | |
| **Neutrophils** | TUM Original | 0.089 ± 0.458 | 0.000 | 0.000 | 0.00 (0.00, 0.00) *ng* | $3.45e^{-2}$ | 0.207 *ns* |
| | Manip to TUM | 0.031 ± 0.124 | 0.000 | 0.000 | | | |
| | NORM Original | 0.045 ± 0.237 | 0.000 | 0.000 | 0.00 (0.00, 0.00) *ng* | $9.70e^{-8}$ | $5.82e^{-7}$ *** |
| | Manip to NORM | 0.104 ± 0.375 | 0.000 | 0.000 | | | |
| **Eosinophils** | TUM Original | 0.056 ± 0.321 | 0.000 | 0.000 | 0.00 (0.00, 0.00) *ng* | 0.582 | 1.00 *ns* |
| | Manip to TUM | 0.027 ± 0.141 | 0.000 | 0.000 | | | |
| | NORM Original | 1.622 ± 2.284 | 0.783 | 2.423 | 0.00 (0.00, 0.11) * | $1.29e^{-8}$ | $7.73e^{-8}$ *** |
| | Manip to NORM | 1.133 ± 2.174 | 0.094 | 1.581 | | | |

**Table S4. Statistical characteristics of patient-level cell-type percentages across groups: original and counterfactual images of the MSI classifier in colorectal cancer.** For each cell type, we report median (%) and interquartile range (IQR, %); mean difference in percentage points (Δ mean) with 95% bootstrap CI; Cliff's δ with 95% bootstrap CI; raw *p*-value (two-sided Mann–Whitney U); and Bonferroni-corrected *p* across six cell types within each block. Percentages were computed per tile and averaged within each patient; the table reports patient-level summaries (median [IQR] across patients). Statistical tests and CIs are computed on the per-patient values. **Block 1:** nonMSIH (Real, *N* = 94) vs. MSIH counterfactuals (CF, *N* = 13). **Block 2:** MSIH (Real, *N* = 13) vs. nonMSIH counterfactuals (CF, *N* = 94). CF, counterfactual; MSIH, microsatellite instability-high.

| Cell Type | Median real [IQR] | Median CF [IQR] | Cliff's δ (95% CI) | Δ mean (pp) 95% CI | *p* raw | *p* corr |
|---|---|---|---|---|---|---|
| **nonMSIH (N = 94 patients) vs. MSIH counterfactuals (N = 13 patients)** | | | | | | |
| **Epithelium** | 63.87 [39.09] | 68.10 [19.59] | 0.155 [-0.108, 0.421] | 9.42 [-0.13, 18.49] | 0.368 | 0.441 |
| **Connective tissue** | 22.88 [28.10] | 18.87 [17.25] | 0.011 [-0.285, 0.316] | -2.09 [-10.16, 6.71] | 0.951 | 0.951 |
| **Plasma cells** | 0.57 [1.63] | 0.29 [0.32] | -0.342 [-0.601, -0.064] | -1.10 [-1.68, -0.52] | 0.047 | 0.155 |
| **Lympho-cytes** | 5.89 [9.82] | 4.03 [7.36] | -0.160 [-0.465, 0.152] | -5.82 [-9.81, -1.77] | 0.353 | 0.441 |
| **Eosinophils** | 0.11 [0.43] | 0.11 [0.12] | -0.200 [-0.444, 0.047] | -0.33 [-0.51, -0.18] | 0.245 | 0.441 |
| **Neutrophils** | 0.04 [0.13] | 0.01 [0.03] | -0.334 [-0.604, -0.050] | -0.09 [-0.16, -0.01] | 0.052 | 0.155 |
| **MSIH (N = 13 patients) vs. nonMSIH counterfactuals (N = 94 patients)** | | | | | | |
| **Epithelium** | 30.70 [42.95] | 48.25 [39.80] | 0.206 [-0.169, 0.568] | 9.86 [-7.30, 26.73] | 0.231 | 0.716 |
| **Connective tissue** | 17.06 [37.72] | 22.41 [27.21] | -0.113 [-0.476, 0.252] | -5.77 [-20.10, 6.89] | 0.514 | 0.716 |
| **Lympho-cytes** | 20.08 [16.34] | 15.83 [15.73] | 0.021 [-0.344, 0.394] | -1.39 [-12.16, 7.73] | 0.905 | 0.905 |
| **Plasma cells** | 3.45 [7.00] | 1.97 [3.13] | -0.092 [-0.499, 0.326] | -1.49 [-4.53, 1.28] | 0.597 | 0.716 |
| **Neutrophils** | 1.75 [2.90] | 0.85 [1.83] | -0.118 [-0.494, 0.273] | -0.97 [-2.66, 0.50] | 0.495 | 0.716 |
| **Eosinophils** | 0.41 [0.94] | 0.28 [0.48] | -0.166 [-0.543, 0.210] | -0.23 [-0.69, 0.14] | 0.336 | 0.716 |

**Table S5. Statistical characteristics of cell type percentages across groups: original and counterfactual images of hepatobiliary cancer types.** For each cell type, the following metrics are reported: mean (%) ± standard deviation (SD); median (%) and the interquartile range (IQR, %); raw *p*-value; corrected *p*-value with Bonferroni correction for multiple comparisons. HCC, hepatocellular carcinoma; CCA, cholangiocarcinoma; Manip to HCC, counterfactual images of CCA (synthetic); manip to CCA, counterfactual images of HCC (synthetic).

| Cell type | Image type | Mean ± SD | Median | IQR | *p* raw | *p* corr |
|---|---|---|---|---|---|---|
| **Epithelial** | HCC Original | 77.928 ± 21.423 | 84.237 | 19.936 | 0.860 | 1.000 |
| | Manip to HCC | 73.425 ± 28.547 | 84.060 | 32.524 | | |
| | CCA Original | 79.001 ± 14.567 | 81.656 | 17.307 | 0.850 | 1.000 |
| | Manip to CCA | 78.411 ± 15.555 | 82.740 | 17.623 | | |
| **Connective** | HCC Original | 14.279 ± 14.870 | 9.234 | 15.134 | 0.456 | 1.000 |
| | Manip to HCC | 12.282 ± 11.738 | 8.824 | 17.576 | | |
| | CCA Original | 15.436 ± 10.646 | 12.770 | 14.344 | 0.495 | 1.000 |
| | Manip to CCA | 14.673 ± 11.698 | 11.067 | 11.980 | | |
| **Lymphocytes** | HCC Original | 6.874 ± 10.281 | 3.795 | 6.171 | 0.804 | 1.000 |
| | Manip to HCC | 12.208 ± 18.863 | 3.049 | 12.428 | | |
| | CCA Original | 5.168 ± 5.016 | 3.615 | 4.237 | 0.254 | 1.000 |
| | Manip to CCA | 6.323 ± 6.234 | 4.315 | 5.724 | | |
| **Plasma cells** | HCC Original | 0.803 ± 2.107 | 0.000 | 0.757 | 0.071 | 0.427 |
| | Manip to HCC | 1.978 ± 4.634 | 0.220 | 1.982 | | |
| | CCA Original | 0.213 ± 0.534 | 0.000 | 0.124 | 0.609 | 1.000 |
| | Manip to CCA | 0.300 ± 0.721 | 0.000 | 0.241 | | |
| **Neutrophils** | HCC Original | 0.087 ± 0.396 | 0.000 | 0.000 | 0.079 | 0.473 |
| | Manip to HCC | 0.105 ± 0.206 | 0.000 | 0.090 | | |
| | CCA Original | 0.148 ± 0.298 | 0.000 | 0.153 | 0.599 | 1.000 |
| | Manip to CCA | 0.213 ± 0.560 | 0.000 | 0.101 | | |
| **Eosinophils** | HCC Original | 0.029 ± 0.184 | 0.000 | 0.000 | 0.541 | 1.000 |
| | Manip to HCC | 0.003 ± 0.016 | 0.000 | 0.000 | | |
| | CCA Original | 0.033 ± 0.109 | 0.000 | 0.000 | 0.362 | 1.000 |
| | Manip to CCA | 0.080 ± 0.301 | 0.000 | 0.000 | | |

**Table S6. Statistical characteristics of cell type percentages across groups: original and counterfactual images of lung cancer types.** For each cell type, the following metrics are reported: mean (%) ± standard deviation (SD); median (%) and the interquartile range (IQR, %); raw *p*-value; corrected *p*-value with Bonferroni correction for multiple comparisons. * indicates significance at $p < 0.001$ after correction. LUAD, lung adenocarcinoma; LUSC, lung squamous cell carcinoma; Manip to LUSC, counterfactual images of LUAD (synthetic); manip to LUAD, counterfactual images of LUSC (synthetic).

| Cell Type | Image type | Mean ± SD | Median | IQR | *p* raw | *p* corr |
|---|---|---|---|---|---|---|
| **Epithelial** | LUAD Original | 73.625 ± 23.669 | 81.255 | 25.964 | $9.646 \cdot 10^{-21}$ | $5.787 \cdot 10^{-2}$[illegible] * |
| | Manip to LUAD | 85.013 ± 16.515 | 91.067 | 16.618 | | |
| | LUSC Original | 67.411 ± 26.451 | 76.779 | 35.111 | 0.154 | 0.922 |
| | Manip to LUSC | 67.617 ± 30.394 | 79.529 | 34.534 | | |
| **Connec-tive tissue** | LUAD Original | 14.737 ± 15.804 | 9.518 | 14.293 | $1.345 \cdot 10^{-24}$ | $8.068 \cdot 10^{-2}$[illegible] * |
| | Manip to LUAD | 7.060 ± 9.201 | 3.661 | 8.578 | | |
| | LUSC Original | 21.770 ± 20.271 | 15.605 | 24.173 | $1.612 \cdot 10^{-3}$ | $9.674 \cdot 10^{-3}$ |
| | Manip to LUSC | 19.315 ± 20.875 | 11.308 | 23.067 | | |
| **Lympho-cytes** | LUAD Original | 9.856 ± 11.023 | 6.675 | 8.695 | $3.167 \cdot 10^{-14}$ | $1.900 \cdot 10^{-1}$[illegible] * |
| | Manip to LUAD | 6.241 ± 8.548 | 3.545 | 6.459 | | |
| | LUSC Original | 8.014 ± 9.821 | 4.840 | 7.970 | $3.761 \cdot 10^{-4}$ | $2.256 \cdot 10^{-3}$ |
| | Manip to LUSC | 10.971 ± 13.553 | 5.997 | 9.555 | | |
| **Neutro-phils** | LUAD Original | 0.275 ± 1.384 | 0.000 | 0.020 | $7.020 \cdot 10^{-9}$ | $4.212 \cdot 10^{-8}$ * |
| | Manip to LUAD | 1.289 ± 4.539 | 0.000 | 0.530 | | |
| | LUSC Original | 1.306 ± 4.683 | 0.000 | 0.582 | $2.095 \cdot 10^{-5}$ | $1.257 \cdot 10^{-4}$ * |
| | Manip to LUSC | 0.228 ± 0.570 | 0.000 | 0.191 | | |
| **Plasma cells** | LUAD Original | 1.378 ± 2.569 | 0.441 | 1.410 | $2.061 \cdot 10^{-28}$ | $1.236 \cdot 10^{-2}$[illegible] * |
| | Manip to LUAD | 0.335 ± 0.812 | 0.000 | 0.249 | | |
| | LUSC Original | 1.343 ± 3.078 | 0.142 | 1.381 | $3.529 \cdot 10^{-8}$ | $2.118 \cdot 10^{-7}$ * |
| | Manip to LUSC | 1.672 ± 2.902 | 0.645 | 2.053 | | |
| **Eosino-phils** | LUAD Original | 0.128 ± 0.495 | 0.000 | 0.000 | 0.019 | 0.113 |
| | Manip to LUAD | 0.062 ± 0.239 | 0.000 | 0.000 | | |
| | LUSC Original | 0.155 ± 0.575 | 0.000 | 0.000 | $7.903 \cdot 10^{-3}$ | $4.742 \cdot 10^{-2}$ |
| | Manip to LUSC | 0.198 ± 0.696 | 0.000 | 0.060 | | |

**Table S7. Statistical characteristics of cell type percentages across groups: original and counterfactual images of breast cancer types.** For each cell type, the following metrics are reported: mean (%) ± standard deviation (SD); median (%) and the interquartile range (IQR, %); raw *p*-value; corrected *p*-value with Bonferroni correction for multiple comparisons. * indicates significance at $p < 0.001$ after correction. IDC, invasive ductal carcinoma; ILC, invasive lobular carcinoma; Manip to IDC, counterfactual images of ILC (synthetic); manip to ILC, counterfactual images of IDC (synthetic).

| Cell Type | Image type | Mean ± SD | Median | IQR | *p* raw | *p* corr |
|---|---|---|---|---|---|---|
| **Epithelial** | IDC Original | 56.829 ± 27.935 | 63.918 | 41.533 | 0.000143 | 0.000859 * |
| | Manip to IDC | 68.565 ± 21.001 | 72.445 | 26.625 | | |
| | ILC Original | 61.331 ± 27.470 | 67.014 | 35.953 | $7.174. \cdot 10^{-42}$ | $4.304 \cdot 10^{-41}$ * |
| | Manip to ILC | 12.481 ± 15.707 | 6.597 | 16.555 | | |
| **Connec-tive tissue** | IDC Original | 34.334 ± 23.086 | 27.084 | 32.571 | 0.000362 | 0.002174 |
| | Manip to IDC | 25.417 ± 18.021 | 22.684 | 22.505 | | |
| | ILC Original | 31.874 ± 24.532 | 24.062 | 33.229 | $4.418 \cdot 10^{-24}$ | $2.651 \cdot 10^{-23}$ * |
| | Manip to ILC | 56.967 ± 15.264 | 57.858 | 18.131 | | |
| **Lympho-cytes** | IDC Original | 7.659 ± 9.367 | 4.460 | 6.310 | 0.010942 | 0.065653 |
| | Manip to IDC | 5.319 ± 6.072 | 3.562 | 5.310 | | |
| | ILC Original | 6.014 ± 7.054 | 3.903 | 5.517 | $2.379 \cdot 10^{-45}$ | $1.428 \cdot 10^{-44}$ * |
| | Manip to ILC | 23.104 ± 13.331 | 20.165 | 17.062 | | |
| **Neutro-phils** | IDC Original | 0.092 ± 0.638 | 0.000 | 0.000 | 0.000308 | 0.001850 |
| | Manip to IDC | 0.052 ± 0.156 | 0.000 | 0.029 | | |
| | ILC Original | 0.015 ± 0.086 | 0.000 | 0.000 | 0.018 | 0.111 |
| | Manip to ILC | 0.060 ± 0.239 | 0.000 | 0.000 | | |
| **Plasma cells** | IDC Original | 1.033 ± 2.543 | 0.357 | 1.081 | 0.000020 | 0.000118 * |
| | Manip to IDC | 0.609 ± 1.326 | 0.087 | 0.486 | | |
| | ILC Original | 0.749 ± 1.717 | 0.200 | 0.718 | $1.713 \cdot 10^{-43}$ | $1.028 \cdot 10^{-42}$ * |
| | Manip to ILC | 7.245 ± 6.354 | 5.765 | 7.005 | | |
| **Eosino-phils** | IDC Original | 0.052 ± 0.230 | 0.000 | 0.000 | 0.547697 | 1.000000 |
| | Manip to IDC | 0.039 ± 0.146 | 0.000 | 0.000 | | |
| | ILC Original | 0.016 ± 0.074 | 0.000 | 0.000 | 0.0796 | 0.478 |
| | Manip to ILC | 0.143 ± 0.625 | 0.000 | 0.000 | | |
